# A PI3K–RAC1 signaling axis downstream of mutant TIE2 promotes endothelial dysfunction in venous malformation

**DOI:** 10.64898/2026.11.12.766208

**Authors:** Komal Sagar, Sandra Schrenk, Chhiring Sherpa, Lauren Crockett, Kara Soroko, Yuqi Cai, Xin Duan, Yi Zheng, Elisa Boscolo

**Affiliations:** Division of Experimental Hematology and Cancer Biology, Cincinnati Children’s Hospital Medical Center, Cincinnati, OH, USA; Department of Pediatrics, University of Cincinnati College of Medicine

**Author notes:** Corresponding author: Elisa Boscolo, Division of Experimental Hematology and Cancer Biology, Cincinnati Children’s Hospital Medical Center, 3333 Burnet Avenue, Cincinnati OH, 45229.

**Keywords:** Venous malformation, TIE2-L914F, RAC1, PI3K signaling, endothelial cells ECs

## Abstract

**Background:** Activating mutations in the endothelial receptor tyrosine kinase TIE2 are a cause of venous malformations (VMs), primarily through dysregulated PI3K/AKT signaling. However, the downstream mechanisms linking constitutively active mutant TIE2 to enlarged vascular lumens, endothelial dysfunction and VM remain incompletely understood. Here, we investigated the role of RAC1 signaling in mutant TIE2 p.L914F driven VM.

**Methods:** Human umbilical vein endothelial cells (HUVECs) expressing wild-type TIE2 or constitutively active mutant TIE2-L914F were analyzed using RAC1 GTP pull-down assays, immunoblotting, and endothelial cell (EC) 2D and 3D functional assays. The role of RAC1 was evaluated using the RAC1 inhibitor NSC23766 in vitro and in vivo xenograft model of VM.

**Results & Conclusions:** ECs expressing mutant TIE2-L914F showed increased active GTP-bound RAC1. TIE2-L914F ECs exhibited cytoskeletal changes with enhanced lamellipodia formation and maturation, accompanied by increased migration. Pharmacological inhibition of RAC1 significantly attenuated the aberrant vascular lumen area and migration in the mutant TIE2 ECs. Mechanistically, we showed that RAC1 hyperactivation is PI3K-dependent but not MEK/ERK-dependent. Consistent with these findings, RAC1 inhibition significantly reduced vascular lesion formation in a xenograft model of VM with TIE2-L914F ECs.

Our study identifies RAC1 as a downstream mediator of TIE2-L914F driven endothelial dysfunction. These results establish a previously unrecognized PI3K-RAC1 signaling axis in the pathogenesis of TIE2-mutated VMs.

## Introduction

Venous malformations (VMs) are the most common type of slow-flow vascular anomaly and are characterized by aberrantly enlarged veins. VMs affect approximately 1 in 5000-10000 individuals and are typically present at birth. Lesions often enlarge progressively throughout life ^1,2^. VM can cause chronic pain, disfigurement, bleeding, and thrombosis resulting in substantial morbidity and severely affecting quality of life ^3,4^. Current treatment options including sclerotherapy and surgical excision are frequently associated with incomplete lesion resolution and high recurrence rates, highlighting the need for targeted molecular therapies ^5–7^. Genetic studies have identified activating mutations in the endothelial receptor tyrosine kinase *TEK* (encoding for TIE2) as the major cause of inherited and sporadic VMs, with TIE2 p.L914F being the most frequent variant ^8–10^. Constitutive TIE2 activation promotes abnormal endothelial cell (EC) signaling, leading to disrupted vascular morphogenesis and lesion formation ^11–13^. In addition to TIE2 mutations, approximately 30% of VM lesions harbor mutations in PIK3CA, the gene encoding the catalytic p110α subunit of phosphoinositide 3-kinase (PI3K) ^14–16^. Previous studies have established that activating TIE2 mutations primarily stimulate the PI3K/AKT signaling pathway, resulting in increased cell survival, proliferation and vascular overgrowth ^12,16,17^. These findings have led to the use of PI3K/mTOR pathway inhibitors, including sirolimus and alpelisib, for the treatment of VMs in affected patients. However, clinical responses often require long-term drug administration while lesions tend to persist, indicating that additional downstream signaling pathways could contribute to the VM pathogenesis. ^18–20^.

During the vasculogenic and angiogenic processes, Rho GTPases function as molecular switches that control vascular lumen formation and maintenance by regulating the activities of the actin cytoskeleton^21,22^. RAC1 activation is necessary for vascular lumen formation during embryonic development and in established three-dimensional (3D) in vitro collagen matrix models ^23–25^. Conversely, RhoA activity is suppressed during EC tubulogenesis ^26–28^.

Although RAC1 is essential for physiological lumen formation^23,25,28^, it is unknown whether excessive or persistent RAC1 activation contributes to the pathological lumen enlargement characteristic of VMs. While recent studies have linked PIK3CA mutations to increased RAC1 activity in vascular malformations, whether a similar mechanism contributes to endothelial dysfunction downstream of mutant TIE2 remains unknown.

In this study, we investigate whether RAC1 hyperactivation functions as a downstream effector of mutant TIE2 signaling and whether it is mediated by increased PI3K signaling. We next determine the effects of RAC1 pharmacological inhibition on the normalization of the endothelial dysfunctions characteristic of VM, such as 3D lumen formation, lamellipodia maturation and increased migration in TIE2-mutant ECs. Lastly, we use an established xenograft model of VM, together with a pharmacological RAC1 inhibitor, to assess its ability to reduce TIE2-mutant EC vascular lesion formation.

## Results

### RAC1 hyperactivation contributes to the ectatic lumen formation in ECs expressing hyperactive mutant TIE2-L914F

We previously reported that HUVEC TIE2-L914F recapitulates the enlarged vascular phenotype characteristic of VM in three-dimensional (3D) fibrin bead assay ^17,29,30^. To investigate the molecular mechanisms underlying the aberrant lumen formation in VM, we first assessed vascular morphogenesis in 3D. As expected, mutant TIE2-L914F ECs formed very enlarged interconnected vascular lumens as compared to HUVECs expressing TIE2-WT and HUVECs non transduced (NT) (Figure 1 A, B and Supplementary figure S1).

**Figure 1:**
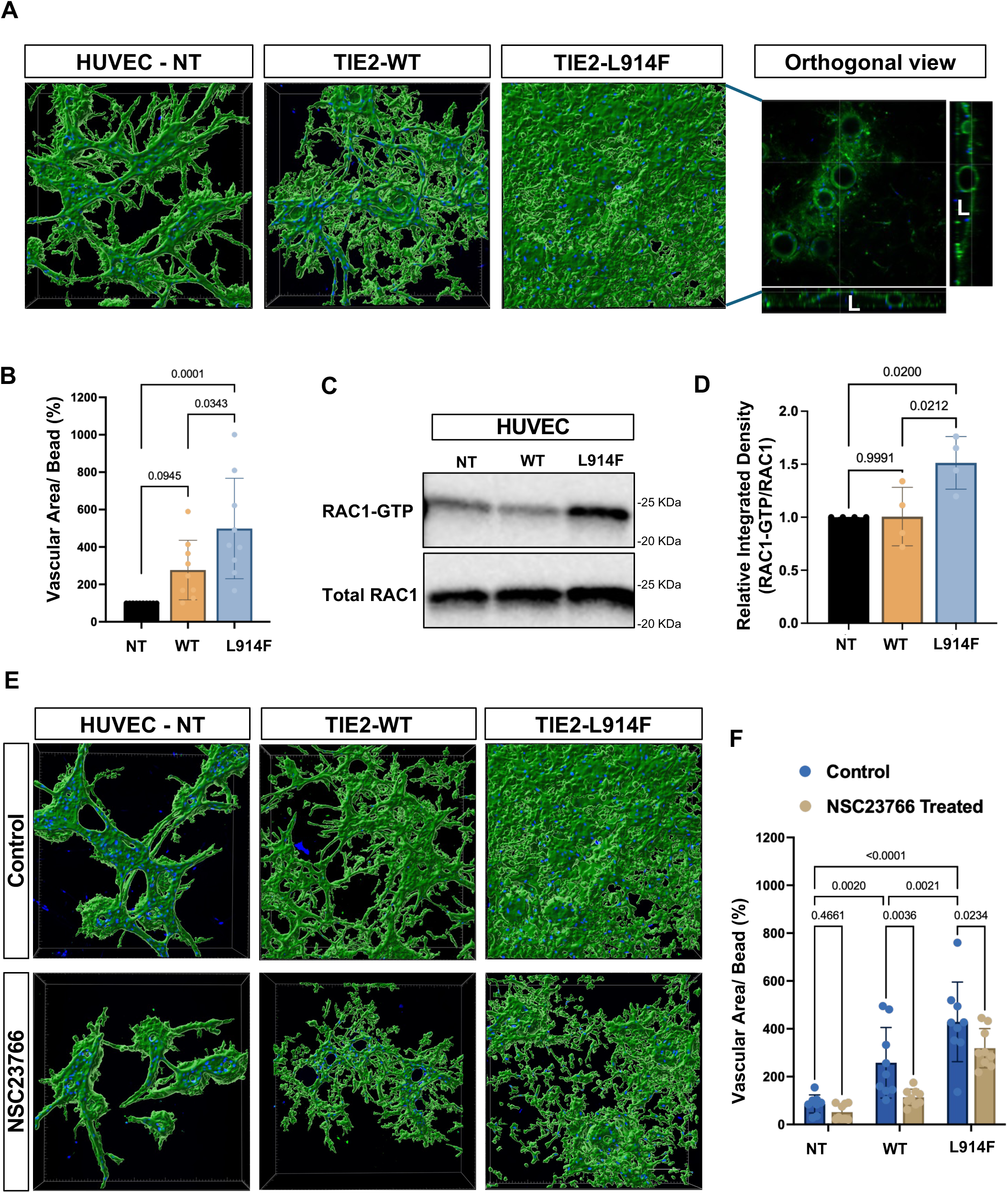
Constitutive TIE2-L914F signaling promotes abnormal vascular morphogenesis through RAC1 activation. **(A)** Representative confocal images of 3D fibrin bead lumen formation assay stained with UEAI (Ulex europaeus agglutinin I, in green – nuclei in blue) shows vascular network formed by non-transduced HUVECs (NT), TIE2-WT, and TIE2-L914F ECs at day 10. L: Lumen. **(B)** Quantification of vascular area normalized per bead number in TIE2-L914F ECs compared with control ECs. **(C)** Immunoblot showing levels of GTP-bound RAC1 and total RAC1 in TIE2-L914F and control HUVECs. **(D)** Quantification of GTP-RAC1 bands normalized to total RAC1 levels. **(E)** Representative confocal images of 3D fibrin bead assay at day 10 of treatment with the selective RAC1 inhibitor NSC23766. **(F)** Quantification of vascular lumen area following NSC23766 treatment or control vehicle, day 10. Data in B, D and F are presented as mean ± SD. Each data point represents one independent experiment (in B, D, F). One-way ANOVA with multiple comparisons (B and D) and 2-way ANOVA with Sidak’s multiple comparison test (F).

Given the central role of RAC1 in lumen formation ^23,25^, we hypothesized that RAC1 hyperactivation contributes to ectatic vein formation in VM. To test this hypothesis, we first investigated RAC1 activation using GTP-RAC1 pull down assay. TIE2-L914F ECs exhibited constitutively increased levels of GTP-bound RAC1 (Figure 1C,D) compared to WT-TIE2 and NT ECs, whereas the total RAC1 transcript and RAC1 protein expression levels remained unchanged (Supplementary figure S2A), indicating that constitutive TIE2 signaling enhances RAC1 activity rather than its expression. Additionally, analysis of CDC42, another important regulator of lumen formation ^27^, showed CDC42 activity was not altered in TIE2-L914F expressing ECs (Supplementary figure S2B).

To determine whether RAC1 activation is functionally required for the VM lumen formation and expansion, TIE2-L914F ECs and control ECs were treated with a selective RAC1 inhibitor, NSC23766 ^31^. To identify concentrations of NSC23766 effective at inhibiting RAC1 activity in L914F ECs we utilized GTP-RAC1 pull-down assay, which demonstrated reduced levels of active RAC1 following 24h treatment with 20μM and 40μM of NSC23766 compared with vehicle-treated control (Supplementary figure S3A). To investigate the role of RAC1 in the formation of aberrant lumens by TIE2-L914F ECs, 3D cultures were treated with 40μM of the RAC1 inhibitor which significantly reduced vascular enlargement in TIE2-L914F ECs while no effect was recorded on proliferation up to 72h (Figure 1E, F, and Supplementary figure S3B). A significant reduction in vascular area was also observed in TIE2-WT ECs following RAC1 inhibition, whereas no significant effect was detected in NT HUVECs.

Collectively, these findings demonstrate that constitutive mutant TIE2-L914F signaling is associated with increased RAC1 activation. Pharmacological inhibition of RAC1 attenuated the abnormal vascular enlargement in a 3D lumen formation assay.

### TIE2-L914F mutation alters lamellipodia and cytoskeletal organization

RAC1 is a master regulator of lamellipodia which consists of dynamic actin-rich membrane protrusions that can form at the cell periphery of the leading edge ^32^.

To analyze lamellipodia organization in the TIE2 mutant EC, we examined the peripheral distribution of membrane F-actin, by classifying ECs based on their protrusive morphology into mature lamellipodia, immature lamellipodia or no lamellipodia (as illustrated in Figure 2A). Based on previous studies ^33,34^, we defined: (i) mature lamellipodia, characterized by continuous thick membrane F-actin-rich protrusions and highly branched F-actin at the cell periphery; (ii) immature lamellipodia, characterized by discontinuous membrane F-actin protrusions; and (iii) absence of lamellipodia, characterized by the lack or very thin F-actin-rich membrane protrusions and lack of peripheral branched F-actin. Quantitative analysis demonstrated that TIE2-L914F ECs showed increased % of cells with mature lamellipodia compared with HUVEC-NT and reduced % of cells with no lamellipodia compared with both TIE2-WT and NT control ECs (Figure 2B and Supplementary figure S4, S5). Treatment with NSC23766 altered lamellipodia distribution in all three endothelial cell groups. In control HUVEC-NT and TIE2-WT ECs, RAC1 inhibition resulted in a slight increase in the proportion of cells lacking lamellipodia (Figure 2C,D). A similar shift was observed in TIE2-L914F ECs, however NSC23766 also induced a 50% reduction in the proportion of cells with mature lamellipodia (Figure 2C,D and Supplementary figure S5). These findings indicate that RAC1 inhibition in TIE2-L914F-expressing ECs promotes a partial normalization of the mutant cell lamellipodia organization. To further investigate the effects of constitutive TIE2 activation on endothelial actin cytoskeletal organization, F-actin content and filopodia formation were quantified by phalloidin staining. TIE2-L914F-expressing HUVECs exhibited a significant increase in phalloidin mean fluorescence intensity (MFI) compared with NT and TIE2-WT cells, indicating enhanced F-actin accumulation (Figure 2C,E). To determine whether these cytoskeletal changes were mediated by RAC1, cells were treated with a RAC1 inhibitor. RAC1 inhibition resulted in about 30% rescue of the F-actin content in the TIE2-L914F ECs (Figure 2E).

**Figure 2:**
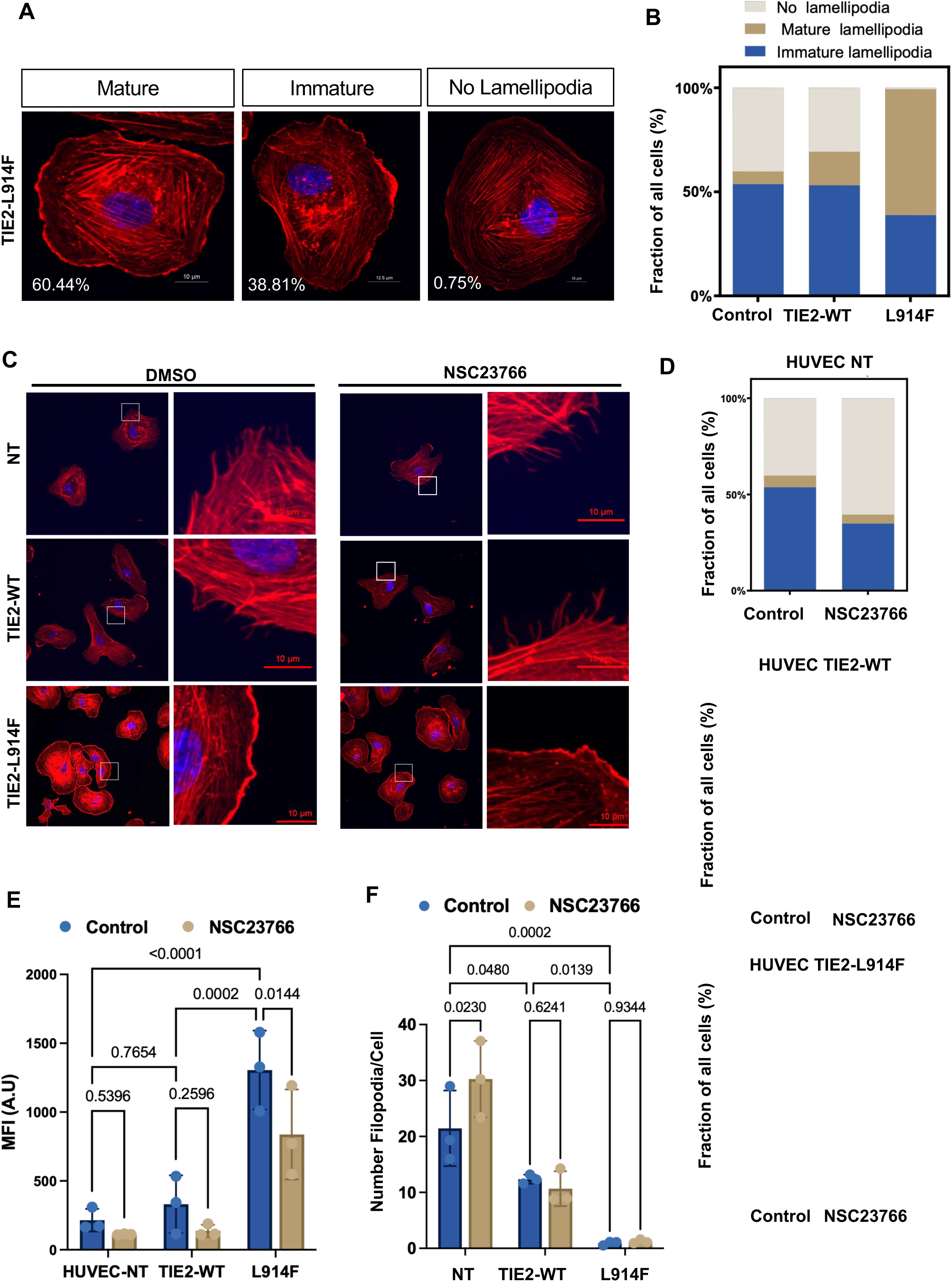
RAC1 inhibition modulates lamellipodia formation in ECs expressing TIE2-L914F. **(A)** Representative confocal images of HUVEC-TIE2-L914F classified according to lamellipodial morphology as mature lamellipodia, immature lamellipodia, or no lamellipodia. F-actin was stained with phalloidin (red) and nuclei with DAPI (blue). Scale bars, 10 μm. **(B)** Quantification of the percentage of cells exhibiting mature lamellipodia, immature lamellipodia, or no lamellipodia in NT, TIE2-WT, and TIE2-L914F HUVEC. **(C)** Representative confocal images of ECs treated with control vehicle or the RAC1 inhibitor NSC23766 and stained for F-actin (phalloidin, red) and nuclei (DAPI, blue). Right images illustrate filopodia in vehicle- and NSC23766-treated cells. Representative images were acquired at 60× magnification. Insets show 2× enlarged views of the selected regions. Higher-magnification insets show selected regions of the cell periphery to facilitate visualization of filopodia. Scale bars, 10 μm. **(D)** Quantification of lamellipodia type in ECs treated with NSC23766. **(E)** Quantification of phalloidin mean fluorescence intensity (MFI), representing cellular F-actin content following RAC1 inhibition with NSC23766. **(F)** Quantification of filopodia number per cell and after NSC23766 treatment. Data are presented as mean ± SD from n=3 independent experiments (B, D, E, F). 2-way ANOVA with Sidak’s multiple comparisons test (E,F). Statistical analysis for B and D is illustrated in Supplementary figure S5.

In contrast, analysis of filopodia revealed a significantly lower number of filopodia in TIE2-L914F ECs compared to both control HUVECs. RAC1 inhibition with NSC23766 did not alter filopodia number in TIE2-WT and TIE2-L914F ECs (Figure 2C high-magnification inset, and 2F). Together, these findings suggest RAC1 hyperactivation contributes to increased lamellipodia maturation and F-actin accumulation in TIE2 mutant ECs, whereas the altered filopodia phenotype appears to be RAC1-independent.

### RAC1 inhibition reduces the migratory capacity of TIE2-L914F ECs

Given the established role of RAC1 in regulating cytoskeletal remodeling, lamellipodium formation and directed migration, we investigated whether RAC1 contributes to the increased migratory phenotype of TIE2-L914F ECs ^29^. TIE2-L914F cells displayed a higher migration rate compared with TIE2 WT and HUVEC NT controls (Figure 3A, B). To determine whether RAC1 contributes to the highly migratory behavior of TIE2-L914F ECs, we performed genetic downregulation of RAC1 by siRNA (Supplementary figure S6). Treatment with RAC1 siRNA delayed wound closure in TIE2-L914F cells, while migration of TIE2-WT ECs was not affected (Figure 3C, D). Consistent with these findings, pharmacological inhibition of RAC1 reduced migration of TIE2-L914F ECs, while migration of NT and TIE2-WT ECs was not affected (Supplementary figure S7). To analyze the migration-specific effects of siRAC1 or NSC23766, the migration experiments were performed for short time, 5 hours, and cell proliferation was inhibited by adding 2 mM hydroxyurea before the start of the experiment. Taken together, these findings demonstrate that the enhanced migratory phenotype of TIE2-L914F ECs is RAC1-dependent.

**Figure 3:**
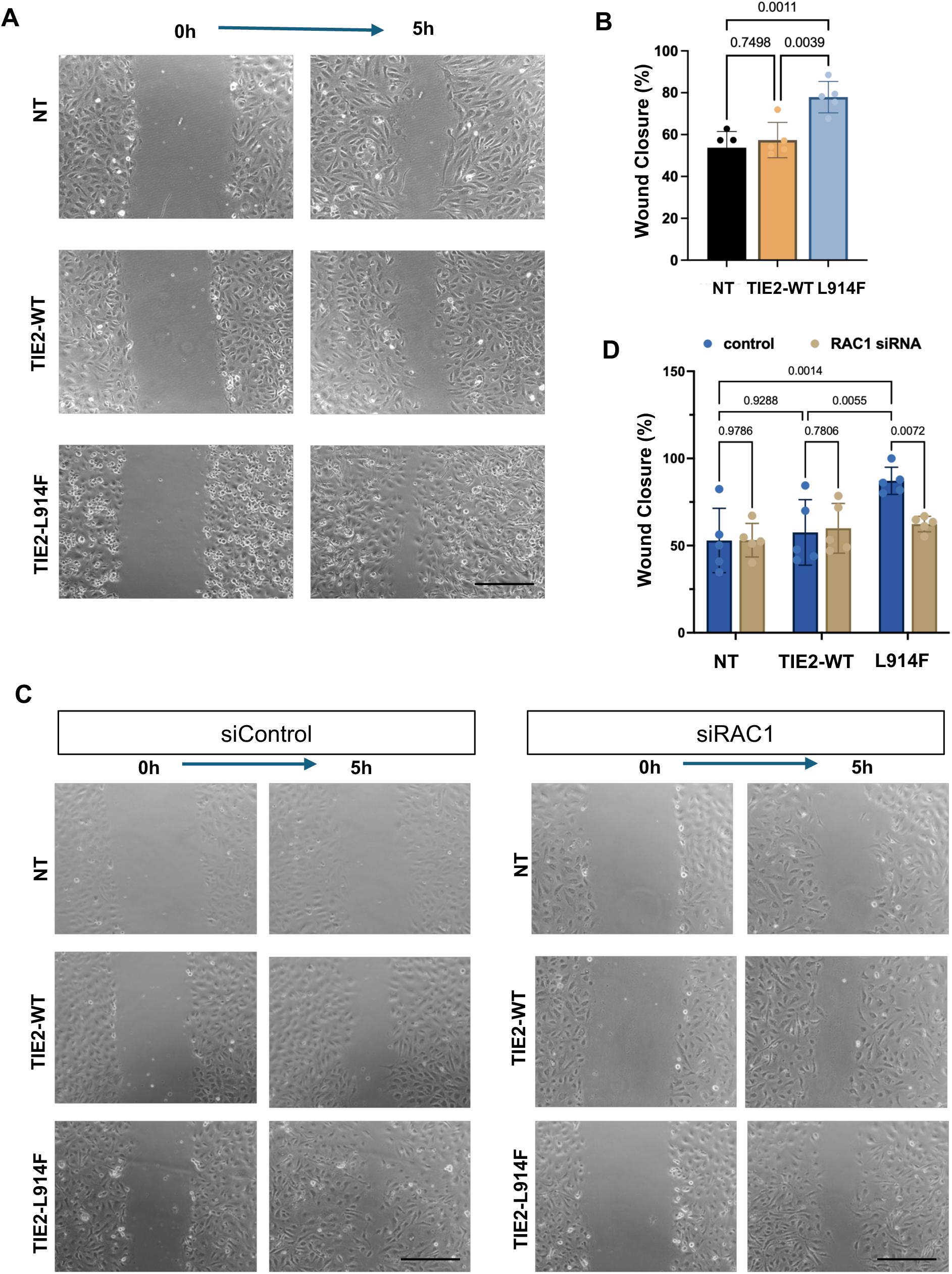
RAC1 regulates EC migration in TIE2-L914F ECs. **(A)** Representative phase-contrast images of scratch wound-healing assays in HUVEC non-transduced (NT), TIE2-WT, and TIE2-L914F ECs at 0 and 5h. Hydroxyurea (2 mM) was included throughout the assay to inhibit cell proliferation. **(B)** Quantification of wound closure at 5 h. n=5 independent experiments. **(C)** Representative phase-contrast images of EC migration following transfection with control siRNA siRNA non-targeting) or RAC1 siRNA in NT, TIE2-WT, and TIE2-L914F ECs. Images were acquired at 0 and 5h. **(D)** Quantification of EC migration following RAC1 siRNA. Data are presented as mean ± SD from n=5 independent experiments. One-way ANOVA with multiple comparisons (B), 2-way ANOVA with Sidak’s multiple comparisons test (D). Scale bar: 200μm.

### PI3K signaling drives RAC1 activation downstream of constitutively active mutant TIE2-L914F

Given that both PI3K/AKT and MAPK/ERK pathways are major downstream effectors of TIE2 signaling ^12,17^, we next sought to determine if these pathways can mediate the hyperactivation of RAC1 observed in TIE2-L914F ECs. To address this, mutant ECs were treated for 2h with the PIK3CA inhibitor, alpelisib or the MEK inhibitor, trametinib, followed by RAC1 pull down assay to assess for RAC1 activation. Alpelisib treatment significantly reduced the levels of GTP bound RAC1 as compared to DMSO-treated TIE2-L914F ECs, indicating that PI3K is required for RAC1 activation (Figure 4 A, B and Supplementary figure S8A,B). In contrast, trametinib had no effect on GTP-RAC1, demonstrating that MAPK/ERK signaling does not contribute to RAC1 activation downstream activation of TIE2-L914F.

**Figure 4.**
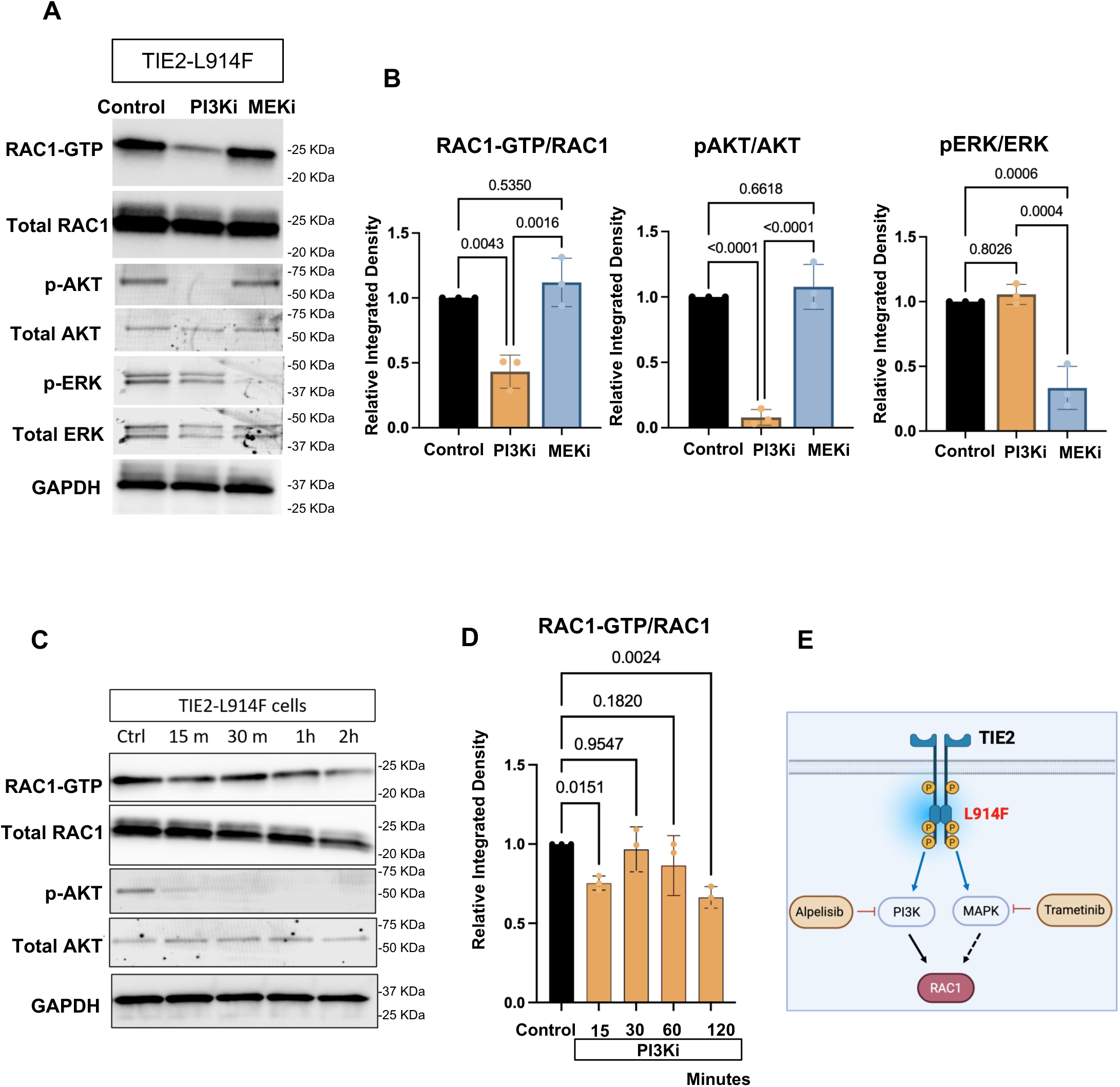
PI3K-dependent regulation of RAC1 activation in TIE2-L914F ECs. **(A)** Representative immunoblots showing RAC1-GTP (pull-down), total RAC1, phosphorylated AKT (p-AKT Ser473), phosphorylated ERK (p-ERK), total AKT, and total ERK in TIE2-L914F ECs treated with the PI3K inhibitor alpelisib, MEK inhibitor trametinib, or control DMSO. **(B)** Densitometric quantification of RAC1-GTP normalized to total RAC1, p-AKT normalized to total AKT, and p-ERK normalized to total ERK following inhibitor treatment. **(C)** Representative immunoblots showing time-course analysis of RAC1 activity following PI3K inhibition with Alpelisib and control DMSO. **(D)** Densitometric quantification of RAC1-GTP normalized to total RAC1 following time-dependent alpelisib treatment. **(E)** Schematic illustration of the proposed signaling mechanism mediated by TIE2-L914F. The L914F mutation activates both the PI3K and MAPK pathways. Pharmacological inhibition of PI3K with alpelisib suppresses PI3K-dependent RAC1 signaling, while inhibition of MAPK signaling did not affect RAC1 activation. Data are presented as mean ± SD from n=3 independent biological replicates. One-way ANOVA with multiple comparisons (B, D).

To establish a time-course of the effect of alpelisib on RAC1 activation, TIE2-L914F ECs were treated for different time durations. Following PI3K inhibition, RAC1 activity progressively decreased over time, with a noticeable reduction at 15 min time point and a more pronounced decrease observed at 2 h post treatment. In parallel, p-AKT was strongly reduced starting at 15 min, confirming the effective inhibition of PI3K signaling while total AKT levels remained unchanged (Figure 4C and 4D). Total RAC1 was also stable across all conditions, indicating that the observed changes were due to altered RAC1 activation rather than total RAC1 protein expression (Supplementary figure S8C).

Together, these results demonstrate that RAC1 hyperactivation downstream of TIE2-L914F is PI3K-dependent, as illustrated in Figure 4E.

### RAC1 inhibition with NSC23766 reduces in vivo lesion formation in a xenograft model of VM

Given the clinical efficacy of PI3K inhibition with alpelisib in patients with *PIK3CA*- and *TEK*-driven vascular malformations ^19,35,36^, and our identification of RAC1 as a downstream effector of PI3K signaling in TIE2-L914F mutant ECs, we set to investigate whether RAC1 inhibition could attenuate the formation of VM-like vascular lesions in vivo.

Mutant TIE2-L914F human ECs were used to perform a xenograft model of VM as previously reported ^11,37^. Upon cell injection, mice were treated daily with the RAC inhibitor NSC23766. A schematic overview of the xenograft experimental design is shown in Figure 5A. Murine body weight was monitored daily to assess for signs of systemic toxicity. We did not record significant weight changes between vehicle control and treated groups (Supplementary figure S9A). Images of explants resected at day 10 (Figure 5B and Supplementary figure S9B), showed that lesions from mice treated with NSC23766 were less vascularized as compared to lesions from vehicle treated controls. Explants from mice treated with NSC23766 also exhibited lower weight compared to vehicle controls (Figure 5C and Supplementary figure S9C). Histological analysis of lesion sections revealed significant reduction in the vascular area (Figure 5D and 5E). Immunostaining with a human-specific lectin, Ulex europaeus agglutinin I (UEA-I), confirmed that blood vessels in vascular lesions consisted of human mutant TIE2-L914F ECs (Figure 5D), distinguishing them from host murine endothelium.

**Figure 5:**
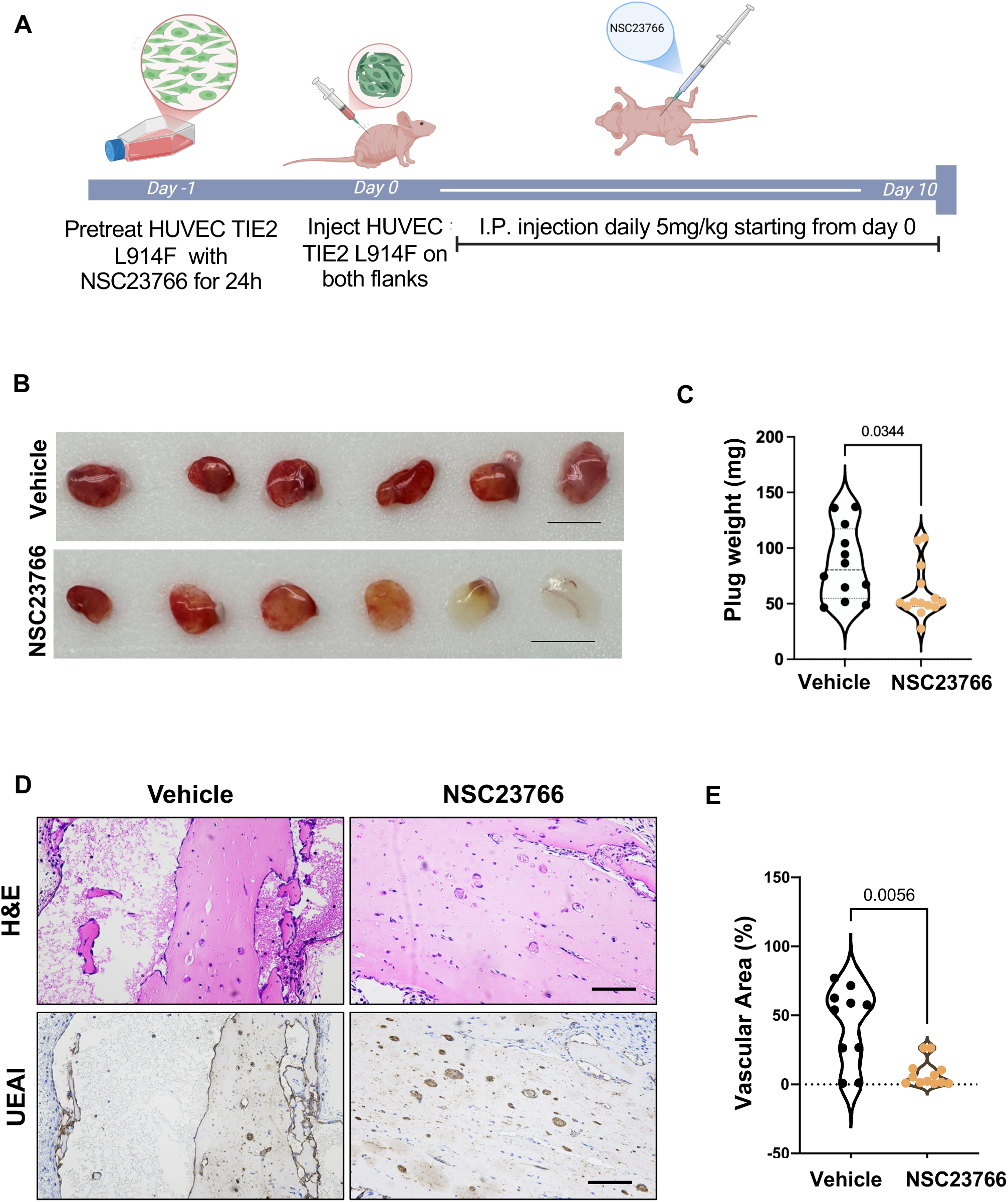
Effect of RAC1 inhibition on TIE2-L914F-driven VM xenografts. (**A**) Experimental schematic of the xenograft study. Human TIE2-L914F ECs were pretreated with NSC23766 for 24 h prior to subcutaneous implantation into both flanks of immunodeficient mice. Mice received daily intraperitoneal injections of NSC23766 (5 mg/kg) or vehicle beginning on the day of implantation and continuing until the study endpoint (Day 10). **(B)** Representative images of xenograft plugs collected from vehicle- and NSC23766-treated mice at the experimental endpoint. Scale bars, 1cm. **(C)** Quantification of xenograft plug weight of mice at the study endpoint. **(D)** Representative images of xenograft sections stained for hematoxylin and eosin (H&E, top panels) and human Ulex europaeus agglutinin-I (UEA-I, bottom panels). Scale bars: 100μm. **(E)** Quantification of vascular area normalized to the explant area is shown. All data points are presented in the violin plot with n=12-14 plugs in each group, with 2 plugs per animal (n=6-7mice/group), Welch’s t-test (C, E).

Together, these findings demonstrate that RAC1 inhibition slows down TIE2-L914F-driven lesion growth in vivo.

## Discussion

In the present study, we identified a mechanistic signaling axis whereby constitutive activation of TIE2-L914F drives pathological features of VM through PI3K-dependent hyperactivation of RAC1. Using complementary in vitro, 3D and in vivo approaches, combined with pharmacological RAC1 inhibition, we demonstrate that RAC1 is hyperactive in TIE2-L914F whereby it contributes to pathological vascular growth. A central observation in this study is that RAC1 hyperactivation in mutant TIE2-L914F ECs contributes to the formation of enlarged vascular lumens in 3D fibrin gel. These findings are consistent with the well-established role of TEK/TIE2 activating mutations as a driver of VM pathogenesis, where aberrant endothelial signaling disrupts vascular organization ^12,17,29,30^. However, while prior studies have largely focused on PI3K/AKT as the principal downstream effector of mutant TIE2, this study significantly extends this paradigm by identifying RAC1 as a downstream effector of TIE2-PI3K in VM.

Our data demonstrate a robust increase in GTP-bound RAC1 in TIE2-L914F ECs without changes in total RAC1 expression, indicating that mutant TIE2 regulates RAC1 primarily through post-translational activation rather than transcriptional or translational control. This is consistent with previous studies showing that stimulation of TIE2 by its ligand angiopoietin-1 (ANGPT-1) preferentially activates RAC1, but not CDC42, to promote EC migration and cytoskeletal remodeling ^38^. Our findings extend these observations by demonstrating that the constitutively active TIE2-L914F mutant induces sustained RAC1 activation in the absence of ligand stimulation, suggesting that persistent RAC1 signaling is a key downstream consequence of pathogenic TIE2 activation. In EC biology, RAC1 is a master regulator of actin polymerization, lamellipodia formation, and directional migration primarily through activation of the WAVE/Arp2/3 complex, and downstream actin branching machinery ^39–41^. Previously, groundbreaking studies by Dr. George Davis showed that inhibition of Rho GTPase by Clostridium toxin B abolishes lumen formation in 3D collagen model ^23,24^ and subsequently identified RAC1 and CDC42 as essential regulators of endothelial lumen formation. Our findings extend these foundational studies by demonstrating that hyperactive RAC1 downstream of mutant TIE2 contributes to excessive vascular enlargement ultimately contributing to the development and progression of VM. Thus, RAC1 appears to function as critical EC regulator as its balanced activation is required for normal vascular development, whereas excessive and persistent activation may contribute to pathological vascular malformation. Pharmacological RAC1 inhibition may also affect normal endothelial morphogenesis in addition to pathological vascular growth. Therefore, dose and treatment duration will need to be carefully evaluated to establish a therapeutic window that limits pathological vascular growth while minimizing effects on normal vascular function.

The enhanced migratory phenotype observed in TIE2-L914F ECs is likely driven by sustained RAC1-dependent lamellipodia formation and actin cytoskeleton remodeling. Lamellipodia are broad, actin-rich membrane protrusions that develop at the leading edge of migrating cells and are essential for directional migration during angiogenesis and vascular morphogenesis ^32,42,43^. RAC1 is the principal regulator of lamellipodia formation through activation of the WAVE regulatory complex and the Arp2/3 complex, which promote branched actin polymerization and membrane protrusion. Under physiological conditions, transient RAC1 activation ensures coordinated EC movement. However, persistent RAC1 activation can enhance protrusive activity and contribute to vascular malformations ^44^. Consistent with this mechanism, TIE2-L914F cells displayed a greater proportion of mature lamellipodia together with enhanced migration, whereas pharmacological inhibition of RAC1 reduced mature lamellipodia and significantly impaired migration. These findings indicate that constitutively active mutant TIE2 signaling promotes pathological EC motility through sustained RAC1 activation and suggest that excessive lamellipodia formation is a key cellular mechanism contributing to the abnormal EC behavior observed in VM.

One important mechanistic finding of this study is the identification of PI3K as the upstream regulator responsible for RAC1 hyperactivation in TIE2-L914F mediated VM. This finding is consistent with previous studies demonstrating that RAC1 is a critical downstream effector in PIK3CA-driven VM ^44^. In that model, RAC1 activation promotes EC migration, cytoskeletal remodeling, and abnormal vascular morphogenesis while pharmacological inhibition of RAC1 reduces lesion development ^44^. Our results broaden this knowledge by demonstrating RAC1 hyperactivation downstream of mutant TIE2 driven VM. The pathological relevance of enhanced RAC1 signaling is further supported by studies in other vascular diseases. Increased RAC1 activity has been reported in varicose veins, where it contributes to endothelial dysfunction and pathological vascular remodeling through increased oxidative stress ^45^. In addition, RAC1 has been shown to regulate endothelial nitric oxide synthase (eNOS) activation and nitric oxide production ^46^ suggesting that sustained RAC1 activation downstream of mutant TIE2 may also enhance eNOS signaling which could promote vasodilation. The pathological relevance of RAC1 extends beyond TIE2-driven VM, as dysregulated RAC1 signaling has been implicated in PIK3CA-driven VM ^44^ and inherited osseous vascular malformation caused by loss-of-function ELMO2 mutations ^47^, suggesting that aberrant RAC1 activation may represent a common mediator of vascular remodeling and endothelial dysfunction.

Here we show that pharmacological inhibition of PI3K with alpelisib effectively reduced GTP-RAC1 whereas in our experimental conditions, inhibition of the MAPK/ERK pathway using trametinib had no detectable effect on RAC1 activity despite effective suppression of ERK. These results clearly position RAC1 activation downstream of PI3K. Mechanistically, PI3K may regulate RAC1 through the generation of phosphatidylinositol-3,4,5-trisphosphate (PIP3), which facilitates the recruitment and activation of RAC1 guanine nucleotide exchange factors (GEFs), promoting GDP-to-GTP exchange on RAC1. Several GEFs, including P-Rex1, Vav family proteins, DOCK family members, and TIAM1, have been implicated in RAC1 regulation^48–51^. However, we did not directly investigate the activation status or functional contribution of these GEFs to RAC1 activation. Future studies involving targeted genetic or pharmacological modulation of candidate GEFs will be necessary to establish their roles in this signaling pathway.

In vivo, RAC1 inhibition significantly reduced xenograft vascular lesion weight and vascular area without significant changes in body weight during the treatment demonstrating that pharmacological targeting of RAC1 reduces lesion growth and suggest it is well tolerated over the treatment period in a preclinical model of VM. Although no significant changes in body weight were observed during treatment, body weight alone is insufficient to establish systemic safety. Therefore, further studies are needed to evaluate the systemic effects of RAC1 inhibition. Furthermore, since NSC23766 administration was initiated at the time of cell implantation, these findings demonstrate attenuation of lesion formation rather than regression of established vascular malformations.

Although the HUVEC-based xenograft model provides a useful approach to investigate the effects of mutant TIE2 signaling and evaluate therapeutic interventions in vivo, it represents an artificial model of VM and does not fully recapitulate the native vascular microenvironment or the complex interactions between endothelial cells and surrounding host tissues. Future studies using genetically engineered mouse models with mosaic expression of TIE2-L914F ^52^ will be important to validate the PI3K-RAC1 signaling mechanism in a more physiologically relevant setting and further assess the therapeutic potential of RAC1 inhibition in VM.

These findings are particularly important given the limited targeted treatment options for VM. While PI3K and mTOR inhibitors have shown partial clinical benefit on vascular malformation ^11,18,19,35,36,53^, our findings provide proof of concept that modulation of RAC1 activity can attenuate key features of the pathological endothelial phenotype driven by mutant TIE2. Further studies should investigate whether combined inhibition of PI3K and RAC1 provides additive or synergistic effects in TIE2-L914F driven VM.

In summary, this study identifies a previously underappreciated PI3K-RAC1 signaling axis as a mediator of TIE2-L914F-driven VM. Our work provides new insights into the molecular mechanism through which mutant TIE2-L914F disrupts EC behavior and contributes to VM development.

### Limitations

This study has some limitations. First, although NSC23766 is a well characterized RAC1 inhibitor ^31^, confirmation using genetic approaches such as genetic deletion or dominant-negative RAC1 expression would further strengthen the causal role of RAC1 in VM pathogenesis. However, such approaches are technically challenging because RAC1 is an essential regulator of EC survival and cytoskeletal dynamics. Complete RAC1 deletion can impair cell growth or induce cell death, whereas prolonged expression of dominant-negative RAC1 may disrupt normal RAC1-dependent cellular functions, thereby complicating the interpretation of experimental results. Second, the effect of NSC23766 on TIE2-WT EC xenograft was not evaluated as in this model these cells do not form vascular lumens. Third, the present study focused exclusively on the TIE2-L914F mutation. Whether other activating TIE2 variants, such as R849W and Y897C, also activate the RAC1 signaling remains unknown. Based on the ability of these mutations to activate PI3K ^12,13^, it would be safe to speculate that this axis is generalizable to other hyperactive TIE2 mutations in VM. Regardless, further studies using different TIE2 mutant variants will be required to determine whether this mechanism represents a common feature of TIE2-driven venous malformation. Finally, the specific RAC1 GEFs responsible for coupling PI3K signaling to RAC1 activation downstream of mutant TIE2 were not investigated and warrant future study.

## Materials and Methods

### Cell culture

Primary human umbilical vein ECs (HUVECs) encoding either wild type TIE2 (TIE2-WT) or constitutively active mutant TIE2-L914F were made by lentiviral transduction and were not immortalized. These cells were generously provided by Dr Lauri Eklund and Dr Miikka Vikkula and were previously described ^12,16^. Cells were maintained under standard endothelial cell culture conditions, and passages 5–6 were used for the experiments to minimize variability associated with prolonged cell culture. Cells were maintained on 1% (w/v) gelatin coated culture plates in EC growth medium prepared from EC basal medium (Lonza, Cat. No. CC-3156) supplemented with EGM-2 SingleQuotsTM (Lonza, Cat. No. CC-4176) and 10% fetal bovine serum (Cytiva HyClone, Cat. No. SH30910.03). Cells were incubated at 37°C with 5% CO_2_ and routinely maintained. All experiments were performed in accordance with institutional biosafety committee guidelines. Alpelisib (Medchem, 501873163) and trametinib (Medchem, HY-10999) were prepared in DMSO (Vehicle).

### 3D lumen formation assay

The 3D fibrin bead assay was performed according to a previously published protocol ^29^. Briefly, Cytodex 3 microcarrier beads (Sigma) were coated with HUVEC NT, HUVEC-TIE2-WT, or HUVEC-TIE2-L914F cells at a density of 400 cells per bead by incubating for 4 h at 37 °C with gentle flicking every 20 min. The following day, coated beads were resuspended in 2 mg/mL fibrinogen (Sigma) containing 0.15 U/mL aprotinin (Sigma) to a final concentration of 500 beads/mL. Thrombin (Sigma) was added, and 0.5 mL of the bead suspension was dispensed into each well of a 24-well plate. The plates were incubated at 37 °C until the fibrin gels had polymerized. ECs were embedded in 3D matrix cultures and imaged by EVOS (Invitrogen) for brightfield images and confocal microscopy (Nikon A1R laser scanning microscope). Vascular area/bead formation was quantified using ImageJ. For experiments evaluating lumen formation with RAC1 inhibitor, the inhibitor was added to both the fibrin gel and culture media throughout the experiment. Media (containing NSC23766, 40 µM) was replaced every other day.

### Western blot analysis

Cells were rinsed twice with ice-cold phosphate-buffered saline (PBS) and lysed in RIPA buffer (Boston Bioproducts BP-115) supplemented with protease and phosphatase inhibitors (Halt protease and phosphatase Single-Use inhibitor cocktail, PI78442). Lysates were maintained on ice for 1 h with intermittent mixing and centrifuged at 12,000 × g for 10 min at 4 °C. Protein concentration was determined using the Pierce 660 nm Protein Assay (Thermo Fisher Scientific, PI23225). Equal amounts of protein were denatured in Laemmli sample buffer by heating at 95 °C for 10 min. Proteins were resolved by electrophoresis on 4-15% or 7.5% Criterion polyacrylamide gel (BioRad) and transferred to PVDF membranes. Membranes were blocked with 5% skimmed milk prepared in TBST for 1 h at room temperature, followed by incubation with primary antibodies overnight at 4 °C (antibodies are listed in Table S1). After washing with TBS, membranes were incubated with horseradish peroxidase (HRP)-conjugated secondary antibodies for 1 h at room temperature. Protein bands were detected using an enhanced chemiluminescence substrate (Millipore Luminata Classico Western HRP substrate, MILL-WBLUC0500) and visualized using a chemiluminescence imaging system. Densitometric analysis of immunoblot signals was performed using ImageJ software, with target protein expression normalized to the corresponding loading control. Antibody information is provided in Supplementary Table S1 and original blots in Supplementary figure S8.

### RAC1 pull-down

RAC1 activity was measured using a PBD-PAK pull-down kit (Thermofisher, 16118) according to the manufacturer’s protocol. Cells were allowed to grow to 80% confluence. Cell lysates were prepared and 1,000 µg of total protein was used for the pull-down assay to isolate the active, GTP-bound form of RAC1. Following affinity precipitation with PAK1-PBD agarose beads, bound proteins were eluted, separated by SDS-PAGE, and analyzed by immunoblotting using an anti-RAC1 antibody. Whole-cell lysates were analyzed in parallel to determine total RAC1 expression.

### Scratch-wound healing assay

For RAC1 siRNA experiments, ECs were seeded in 24-well plates at a density of 2 × 10 cells/cm² and grown until a uniform confluent monolayer was formed. A straight wound was generated across the monolayer using a sterile 20-µL pipette tip. Bright-field images of the wound were acquired immediately after scratching (0 h and subsequently at 20-min intervals for up to 5 h using an EVOS imaging system (Invitrogen). Wound closure was quantified using Fiji (ImageJ) by measuring the remaining cell-free area at each time point and expressing the results as the percentage of the original wound area at 0 h.

For NSC23766 (MedChem, HY-15723A, prepared in sterile water, vehicle) experiments, two-chamber culture insert system (Ibidi, Cat. No. 80209) was used to generate a defined cell-free gap of approximately 500 µm in glass-bottom 24-well plates (Cellvis, Cat. No. P24-1.5H-N). A total of 2,000 cells per chamber were seeded and allowed to form confluent monolayers. To minimize the contribution of cell proliferation, the medium was supplemented with 2 mM hydroxyurea throughout the assay. Migration was initiated by removal of the silicone insert, and cells were imaged live at 20 min intervals for up to 5 h using a Nikon Ti2 widefield microscope maintained at 37 °C and 5% CO₂. The NSC23766 inhibitor was included throughout the assay.

### siRNA-Mediated Gene Knockdown

Gene knockdown was carried out using siRNA transfection with DharmaFECT transfection reagent (T-2004-02) (siRAC1: AUGAAAGUGUCACGGGUAA; siNT, non-targeting: UAGCGACUAAACACAUCAA) reagent according to manufacturer’s protocol. Briefly, cells were seeded in antibiotic-free complete culture medium and allowed to grow overnight under standard growth conditions. For transfection, 50 nM of siRNA and DharmaFECT™ reagent were each diluted in serum free media and incubated for 5 min at RT. Then they were combined and incubated for 20 min to allow formation of siRNA-lipid complexes. Cells were either transfected with RAC1 siRNA or non-targeting control siRNA (siNT) for 48-72h before analyses.

### **q-** PCR

Total RNA was isolated from cells using an RNA extraction kit (Qiagen, cat 74106) according to the manufacturer’s instructions. RNA concentration and purity were determined using nanodrop. cDNA was prepared using reverse transcription kit (Bio-Rad, cat#1708841). qPCR was performed using SYBR Green master mix on a Biorad CFX96 system. Relative gene expression was calculated after the normalization to the housekeeping genes Hypoxanthine phosphoribosyl transferase 1 (*HPRT1*) and TATA-binding protein 1 (*TBP1*). For each gene, a relative standard curve was generated to assess amplification performance. Primer amplification efficiencies greater than 90% were considered acceptable All the reactions were performed in triplicate, and results are presented as relative fold change compared with the control group. Primer sequences are RAC1 Forward: 5’ TACGCCCCCTATCCTATCCG 3’ RAC1 Reverse: 5’ CAATCGGCTTGTCTTTGCCC 3’, HPRT Forward: 5’ CCT GGC GTC GTG ATT AGT GAT 3’ HPRT Reverse: 5’ GGG CTA CAA TGT GAT GGC CT 3’, TBP1 Forward: 5’ GTG GGG AGC TGT GAT GTG AA 3’, TBP1 Reverse: 5’ TGC TCT GAC TTT AGC ACC TGT 3’

### Immunofluorescence

Cells were seeded onto gelatin coated coverslip and allowed to adhere before fixation with 4% paraformaldehyde for 20 min at RT. Following fixation cells were blocked with 3% BSA containing 0.1% Triton X-100 for 1 h at RT. To visualize the actin cytoskeleton, samples were stained with rhodamine-conjugated phalloidin (Life technologies, R415) for 30min at RT. Cell nuclei were counterstained with DAPI (4′,6-diamidino-2-phenylindole) stain for 30 min at RT. Confocal images were captured using Nikon A1R inverted confocal microscope. To categorize cells for lamellipodia formation, cells were randomly selected and analyzed based on the organization of the peripheral F-actin network. Cells lacking a peripheral actin-rich protrusion were classified as having no lamellipodia. Cells with small, discontinuous, or ruffled actin-rich protrusions at the cell edge were classified as immature lamellipodia. Cells exhibiting broad, continuous and well-organized actin-rich membrane protrusions were classified as having mature lamellipodia. Images were quantified using NIS elements for lamellipodia classification.

### Mouse Xenograft model

All animal procedures complied with ARRIVE guidelines and were conducted in accordance with relevant guidelines and regulations, including the ‘Guide for the Care and Use of Laboratory Animals’. Procedures were approved by the Institutional Animal Care and Use Committee of Cincinnati Children’s Hospital and Medical Center (IACUC No: 2026-0023), according to the NIH Guide for the Care and Use of Laboratory Animals.

Human umbilical vein ECs (HUVECs) encoding either wild type TIE2 (TIE2-WT) or constitutively active mutant TIE2-L914F were expanded in culture and where indicated were treated with NSC23766 at 40μM for 24 h before the implantation of cells. Three mice received TIE2-WT cells, six mice received TIE2-L914F cells with vehicle treatment, and seven mice received TIE2-L914F cells with NSC23766 treatment. A total of 2.0 × 10^6 cells were resuspended in 200 µL Matrigel (CB40234C, Corning) and injected subcutaneously into both flanks of 6–7-week-old male athymic nude (nu/nu) mice (Envigo). Mice were treated daily with NSC23766 prepared in PBS (Vehicle) and administered by intraperitoneal (i.p.) injection at a dose of 5 mg/kg. After 10 days, Matrigel plugs were harvested, weighed, fixed in 4% paraformaldehyde, and embedded in paraffin. Sections (5 µm) were stained with hematoxylin and eosin (H&E). Vascularization was evaluated by capturing five random fields per section using Nikon Eclipse Ci and representative images were taken from Agilent lionheart Imaging system. Vascular area was quantified as the percentage of vessel-containing area relative to total field area using ImageJ software.

All experiments were carried out in accordance with institutional biosafety and chemical safety guidelines and regulations.

### Proliferation assay

EC proliferation was assessed using live cell imaging. 3500 cells were seeded on to each well of 96 well plate coated with 1% gelatin. Cells were stained with NucBlue (1 drops/20 mL, Invitrogen, R37605). Cells were imaged every 2 h for 72h using a Lionheart FX Live Cell Imaging System (Agilent). NucBlue positive cells were counted using Gen5, Lionheart software.

### Statistical analysis

Data was represented as mean±SD from at least three biological experiments, with exact number of replicates provided in the figure legend. Differences between two groups were analyzed using an unpaired t-test. For more than 2 groups one way or two-way ANOVA with post hoc test was applied. All statistical analyses were performed using GraphPad Prism (Version 10). A P value of <0.05 was considered statistically significant.

## Acknowledgements and sources of funding

Research reported in this manuscript was supported by the National Heart, Lung, and Blood Institute, under Award Numbers R01HL117952 (E.B.), R01HL167700 (E.B.), and 5R25CA261610-05 (PI: M.S. Butschkovacic, supporting L.C.), part of the National Institutes of Health. The content is solely the responsibility of the authors and does not necessarily represent the official views of the National Institutes of Health. This manuscript is the result of funding in whole or in part by the National Institutes of Health (NIH). It is subject to the NIH Public Access Policy. Through acceptance of this federal funding, NIH has been given a right to make this manuscript publicly available in PubMed Central upon the Official Date of Publication, as defined by NIH. We thank the Bio-imaging and Analysis Facility and the Comparative Medicine Division at Cincinnati Children’s Hospital Medical Center for providing state-of-the-art instrumentation, services, training, and education.

## Author contributions

E.B., K. Sagar. and S.S. and conceived the project. E.B. supervised the research. Y.C., X.D., Y.Z. assisted with initial investigations on RAC1 and use of NSC23766 in vitro. K. Sagar., C.S., K. Soroko. and L.C. performed the mouse experiments. K. Sagar performed in vitro experiments, immunostaining and confocal imaging. K. Sagar wrote the initial manuscript and figure draft. All authors revised and commented on the manuscript.

## Data availability statement

All data generated or analyzed during this study are included in this published article and its Supplementary Information file.

## Abbreviations

EC: endothelial cells
VM: venous malformation
HUVEC: human umbilical vein EC
TEK (TIE2): tyrosine kinase with immunoglobulin-like and EGF-like domains 2
F-actin: filamentous actin

