## Supplementary Data for "A PI3K–RAC1 signaling axis downstream of mutant TIE2 promotes endothelial dysfunction in venous malformation"

Content:

Supplementary Figures S1-S10

Supplementary Table S1

#### Supplementary Figure S1

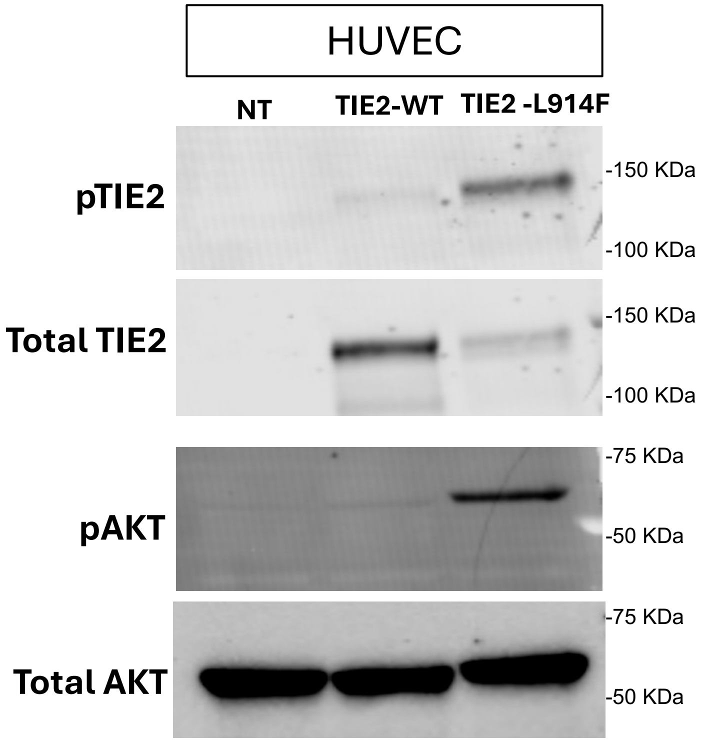


**Supplementary Figure S1 (For data in Figure 1): HUVEC-TIE2-L914F show constitutive TIE2 and AKT hyperactivation.** Immunoblot for phosphoTIE2, phosphoAKT and respective total protein levels, in HUVEC Non-Transfected (NT) HUVEC-TIE2-WT (Wild-type) and HUVEC-TIE2-L914F.

#### Supplementary Figure S2

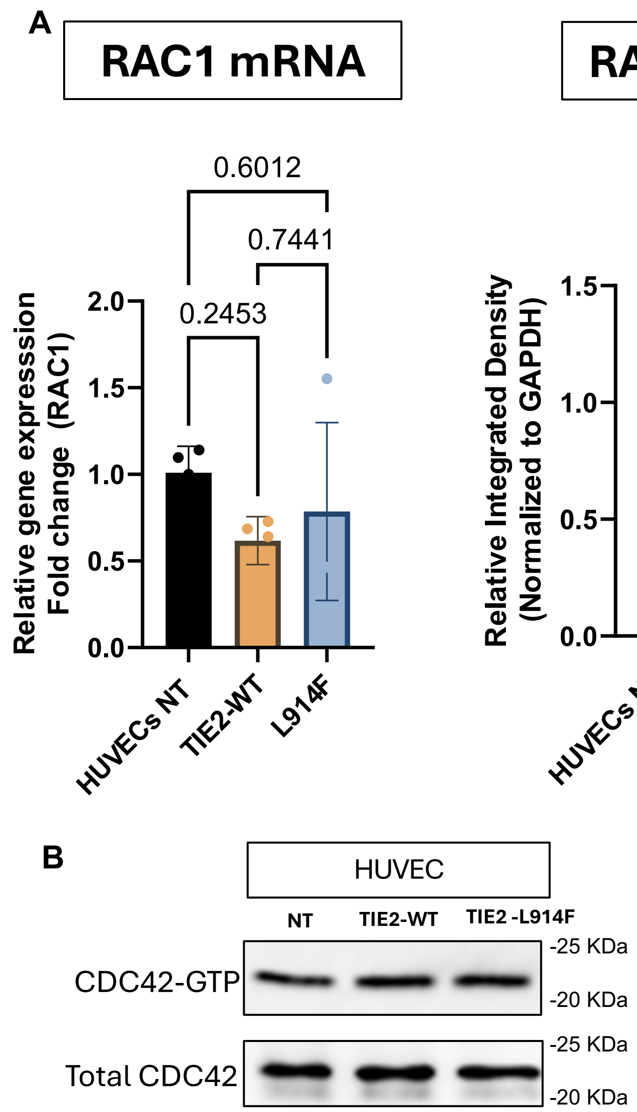


**Supplementary Figure S2 (For data in Figure 1): HUVEC-TIE2-L914F show increased RAC1 activation but not CDC42 activation - with no changes in RAC1 total levels**. Quantification of **(A)** RAC1 mRNA and total RAC1 protein expression in HUVECs NT, TIE2-WT and TIE2-L914F cells. **(B)** Representative CDC42-GTP pull-down assay and corresponding input (total CDC42).

#### Supplementary Figure S3

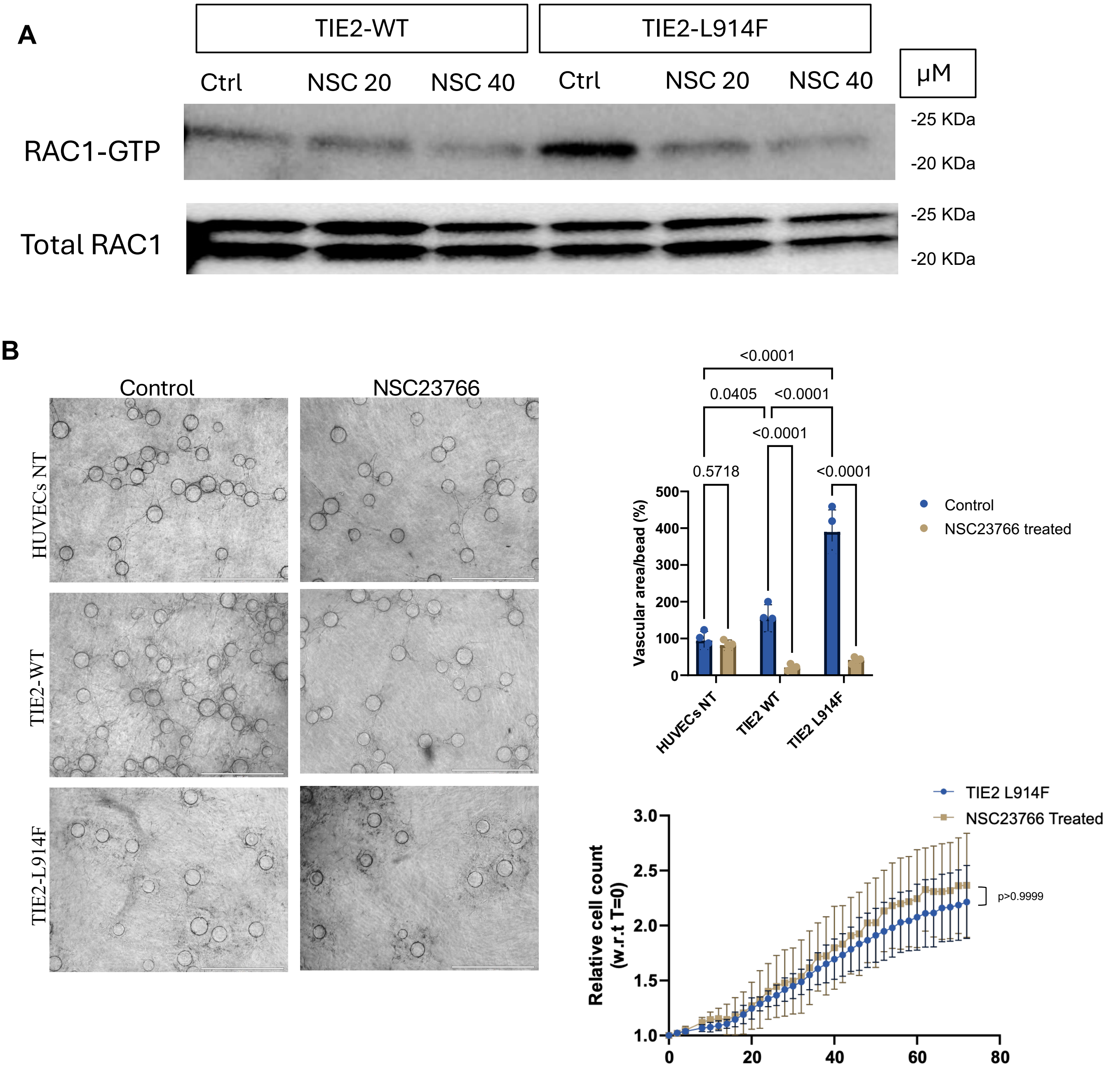


**Supplementary Figure S3 (For Figure 1): RAC1 inhibitor NSC23766 effects in vitro and in 3D lumen formation assay. (A)** Representative immunoblot showing active RAC1 (RAC1-GTP) after 24 h of NSC23766 treatment. Total RAC1 in whole-cell lysates is shown as the loading control. NSC23766 (40 μM) treatment decreased RAC1-GTP levels compared with the vehicle-treated control, indicating effective inhibition of RAC1 activity. **(B)** Representative bright-field (BF) images of 3D lumen-forming cultures with vehicle or NSC23766. Scale bar: 100μm. Quantification of vascular lumen area from phase images. n=3 independent experiments. Data presented as mean ± SD. 2-way ANOVA with Sidak's multiple comparisons test. Bottom left graph illustrates cell proliferation curves for HUVEC-TIE2 L914F following 40μM NSC23766 treatment. n=4 technical replicates. 2-way Anova

#### Supplementary Figure S4

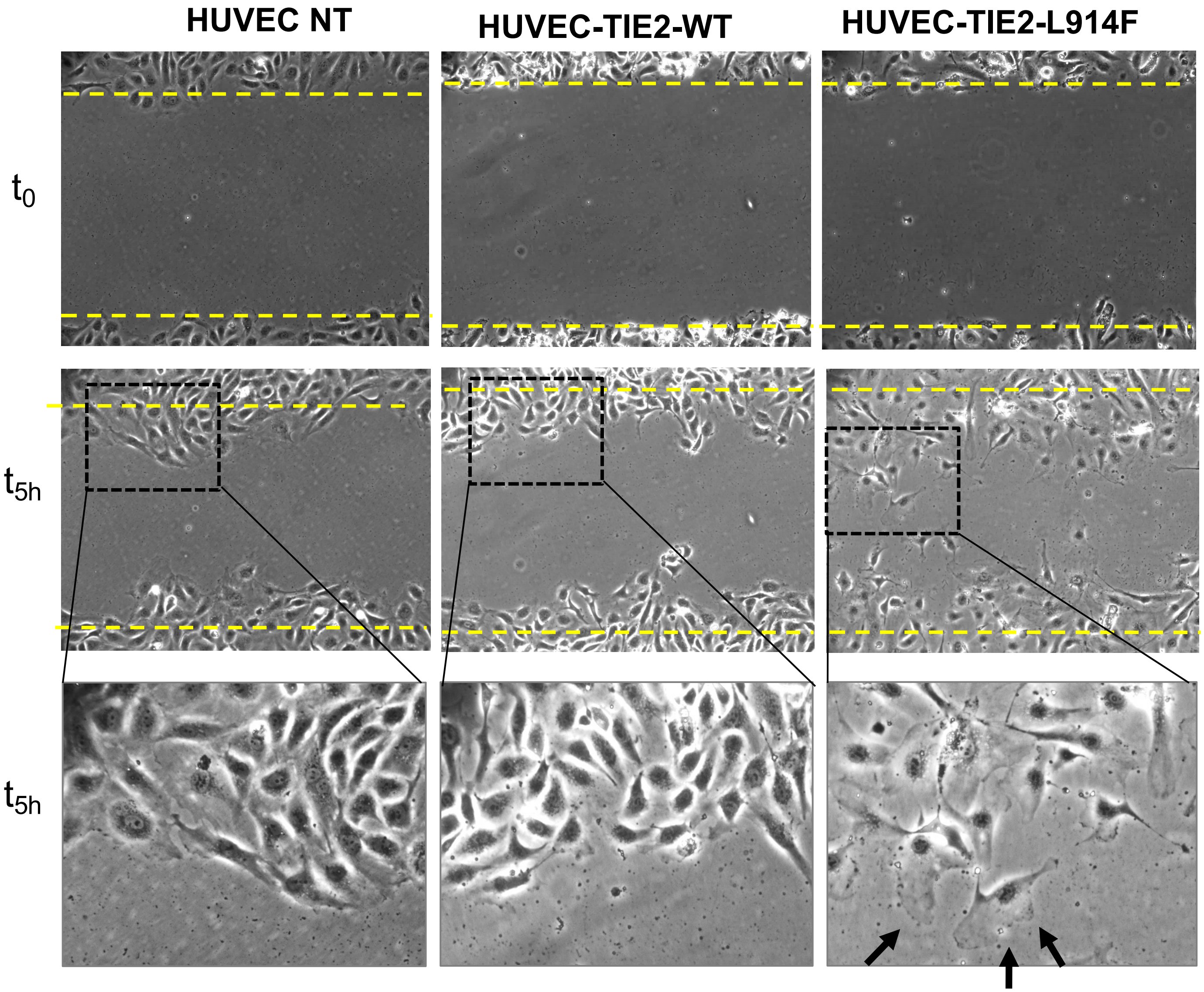


**Supplementary Figure S4 (For Figure 2): Lamellipodia formation in TIE2 mutant EC during migration.** HUVEC NT, HUVEC TIE2-WT and HUVEC-TIE2-L914F were subjected to migration upon performing scratch wound and imaging performed right after the scratch was made (t0) and after 5h. Insets show high magnification of migrating front and black arrows point to lamellipodia.

### Supplementary Figure S5

**A**

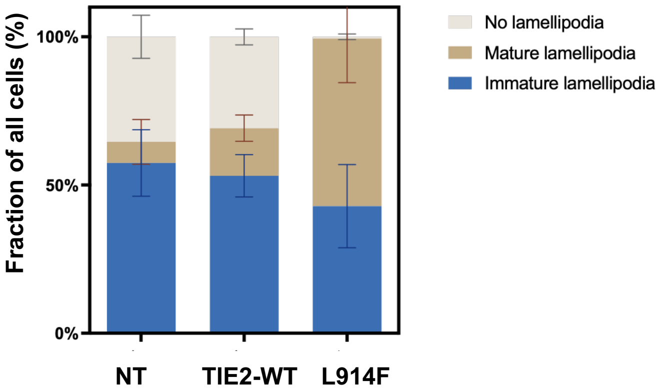


| Tukey's multiple comparisons test | Summary | Adjusted P Value |
|---|---|---|
| Immature lamellipodia | | |
| HUVECs NT vs. TIE2 WT | ns | 0.8452 |
| HUVECs NT vs. TIE2 L914F | ns | 0.4237 |
| TIE2 WT vs. TIE2 L914F | ns | 0.5652 |
| Mature lamellipodia | | |
| HUVECs NT vs. TIE2 WT | ns | 0.3076 |
| HUVECs NT vs. TIE2 L914F | * | 0.0301 |
| TIE2 WT vs. TIE2 L914F | ns | 0.0635 |
| No lamellipodia | | |
| HUVECs NT vs. TIE2 WT | ns | 0.6150 |
| HUVECs NT vs. TIE2 L914F | * | 0.0238 |
| TIE2 WT vs. TIE2 L914F | ** | 0.0020 |

**B**

**HUVEC NT**

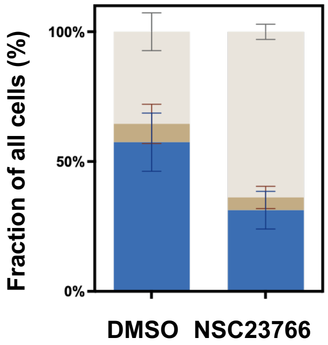


| HUVEC-NT | P value | Mean of DMSO | Mean of NSC23766 Treated | Difference |
|---|---|---|---|---|
| Immature lamellipodia | 0.027172 | 57.46 | 31.28 | 26.18 |
| Mature lamellipodia | 0.689169 | 7.083 | 4.929 | 2.154 |
| No lamellipodia | 0.003305 | 35.46 | 63.79 | -28.34 |

**HUVEC TIE2-WT**

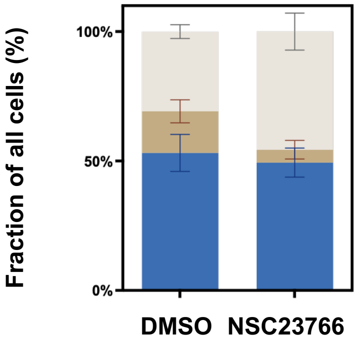


| HUVEC-WT | P value | Mean of DMSO | Mean of NSC23766 Treated | Difference |
|---|---|---|---|---|
| Immature lamellipodia | 0.514644 | 53.12 | 49.38 | 3.742 |
| Mature lamellipodia | 0.028351 | 16.06 | 4.976 | 11.09 |
| No lamellipodia | 0.028209 | 30.82 | 45.65 | -14.83 |

**HUVEC TIE2-L914F**

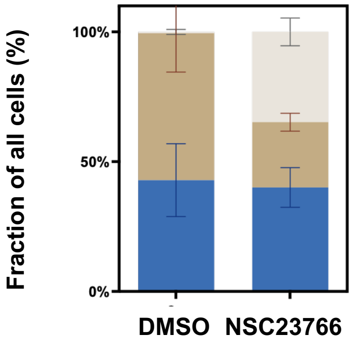


| HUVEC-TIE2-L914F | P value | Mean of DMSO | Mean of NSC23766 Treated | Difference |
|---|---|---|---|---|
| Immature lamellipodia | 0.775208 | 42.89 | 40.08 | 2.817 |
| Mature lamellipodia | 0.023772 | 56.55 | 25.12 | 31.43 |
| No lamellipodia | 0.000402 | 0.5556 | 34.81 | -34.25 |

**Supplementary Figure S5 (For Figure 2): RAC1 inhibition modulates lamellipodia formation in ECs. (A)** Quantification of the percentage of cells exhibiting mature, immature, or no lamellipodia in NT, TIE2-WT, and TIE2-L914F HUVEC. 2-way ANOVA with Tukey's multiple comparisons test. **(B)** Quantification of lamellipodia type in ECs treated with NSC23766. Multiple unpaired T-tests. Data in A and B are presented as mean ± SD from n=3 independent experiments.

#### Supplementary Figure S6

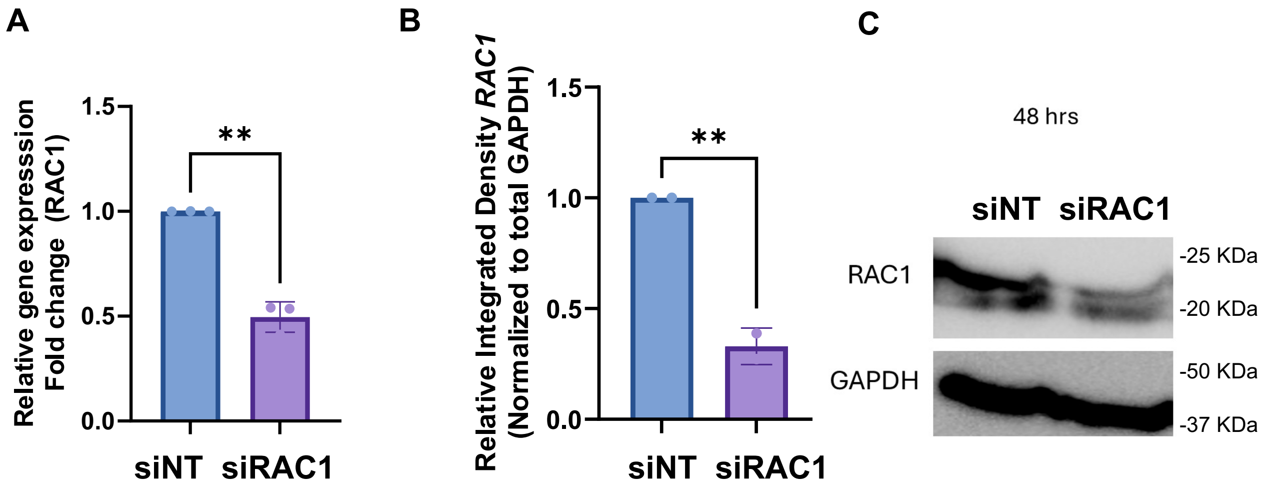


**Supplementary Figure S6 (for data in Figure 3). RAC1 expression in cells treated with short-interference si RNA.** TIE2-L914F endothelial cells were transfected with either non-targeting control small interfering (si)RNA (siNT) or RAC1 siRNA (siRAC1) and analyzed 48 h post-transfection. **(A)** Quantitative RT-PCR analysis showing a significant reduction in RAC1 mRNA expression following siRAC1 transfection compared with the siNT, control. **(B)** Densitometric quantification of RAC1 protein levels normalized to GAPDH demonstrates efficient RAC1 knockdown. (**C)** Representative Western blot showing RAC1 protein expression in siNT- and siRAC1-transfected cells, with GAPDH used as a loading control. All data are presented as mean ± SD from n=3 independent experiments. A and B Welch's t-test. **p<0.001

#### Supplementary Figure S7

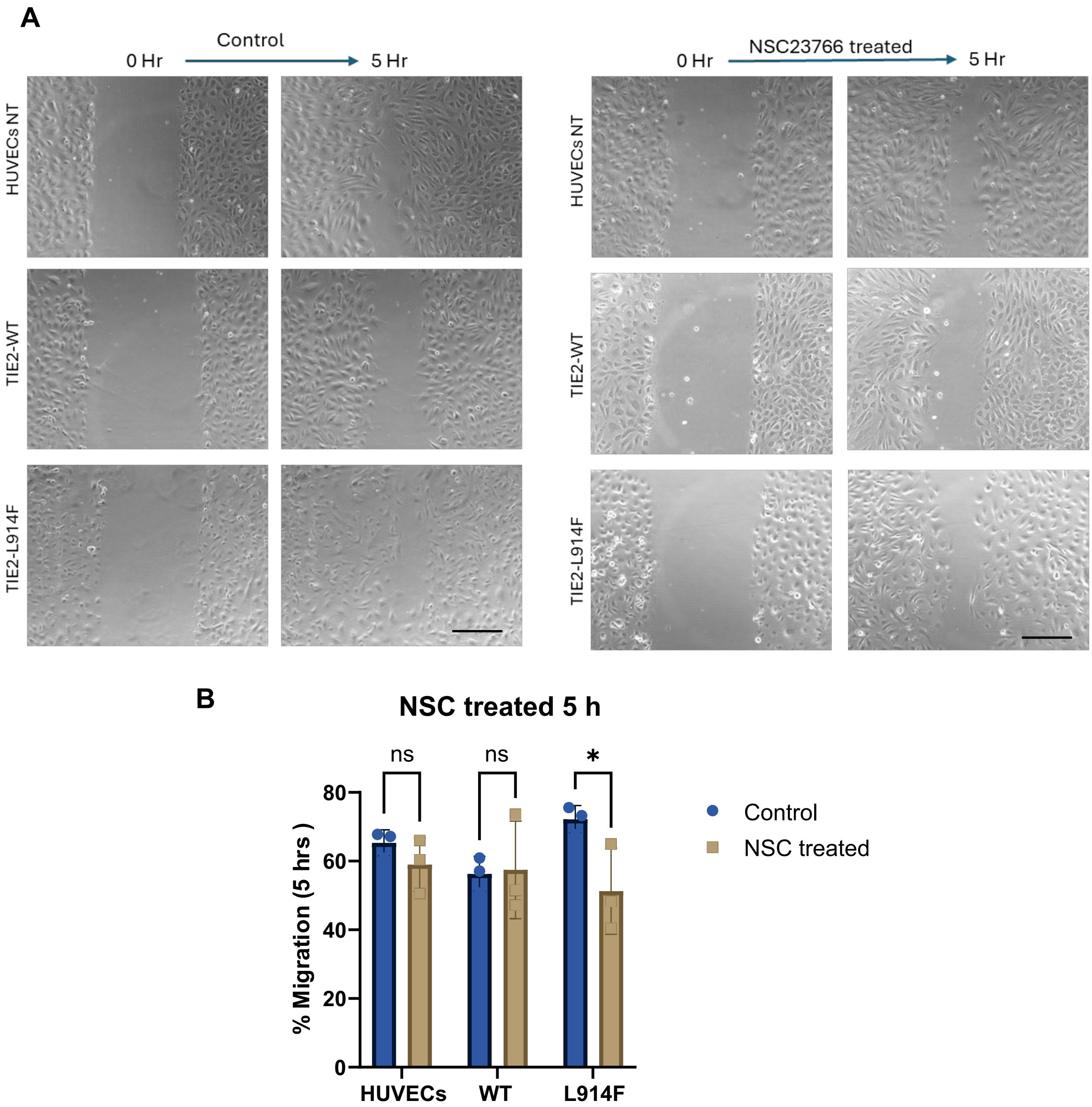


**Supplementary Figure S7 (for data in Figure3): RAC1 inhibition with NSC23766 reduces migration in HUVEC-L914F. (A)** Representative phase-contrast images of wound healing assays. Uniform scratch was introduced using culture inserts into confluent cell monolayers, and wound closure was monitored over 5h time points. Cells treated with the RAC1 inhibitor NSC23766 exhibited reduced wound closure compared with vehicle-treated controls, indicating impaired migratory capacity. Scale bar: 200µm. **(B)** Quantification of wound closure, expressed as the percentage of the initial wound area remaining (or percentage wound closure) at the indicated time points. Data are presented as mean ± SD from n = 3 independent experiments. 2-way ANOVA with Sidak's multiple comparisons test. $^{*}p<0.05$.

#### Supplementary Figure S8

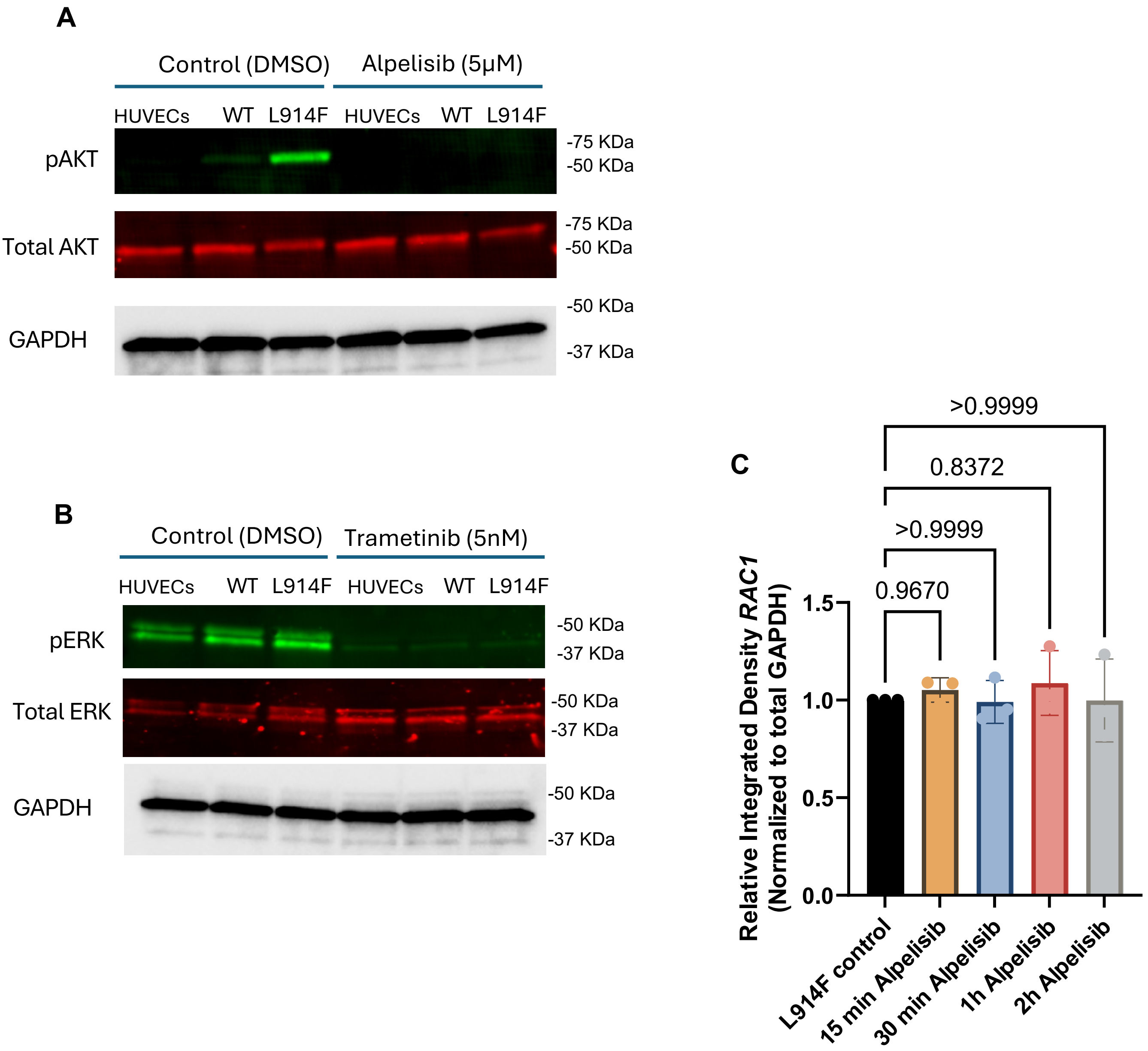


**Supplementary Figure S8 (for data in Figure 4): Treatment with alpelisib rescues increased RAC1 activation in HUVEC-TIE2-L914F.** Representative immunoblot of **(A)** phosphorylated AKT (p-AKT), **(B)** phosphorylated ERK (p-ERK), and the loading control following treatment with alpelisib or trametinib. **(C)** Quantification of total RAC1protein levels normalized by GAPDH. n=3 independent experiments. One-way ANOVA with multiple comparisons.

#### Supplementary Figure S9

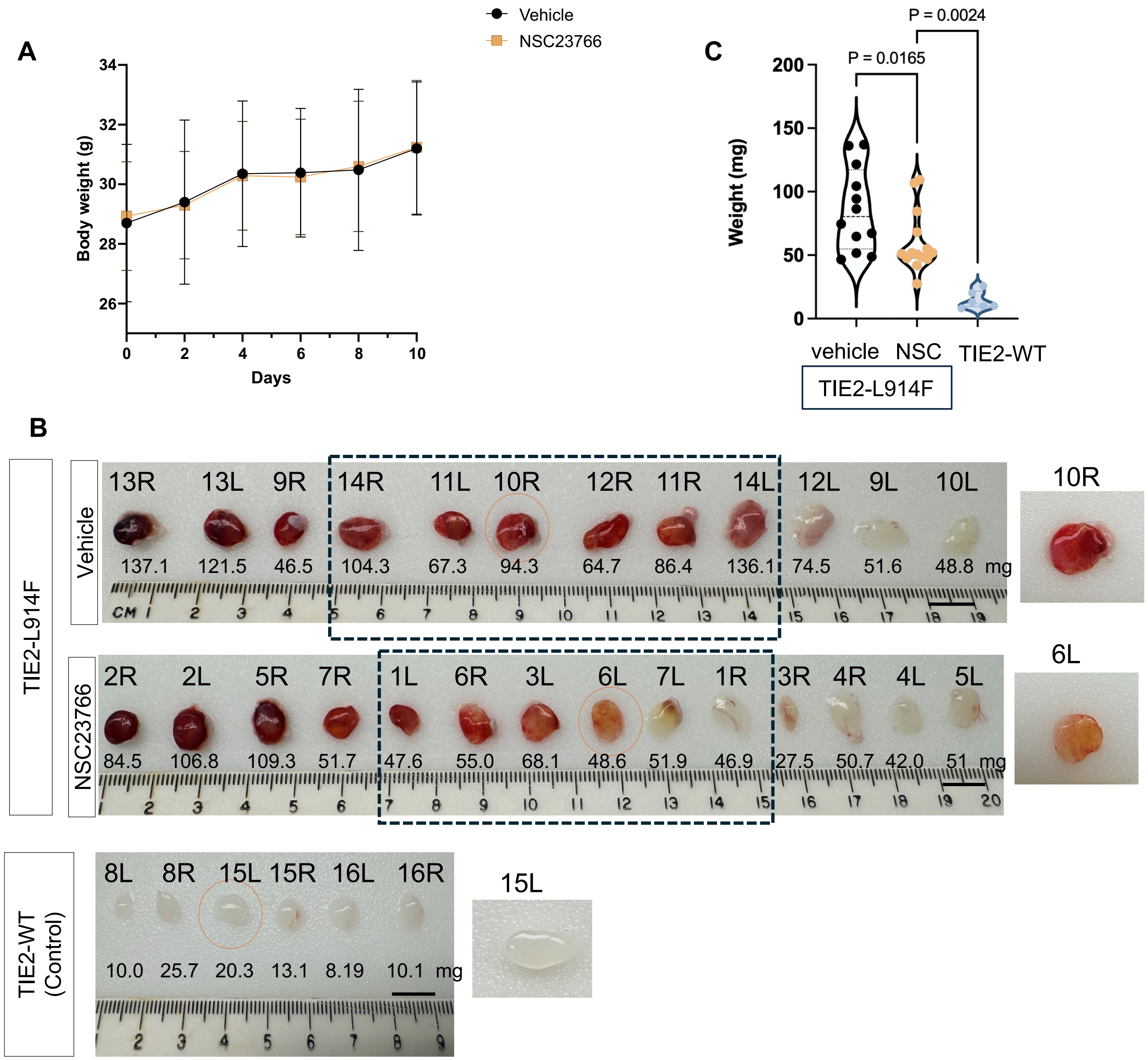


**Supplementary Figure S9 (for data in Figure 5): RAC1 inhibition reduces VM xenograft lesion vascularity without toxicity. (A)** Mouse body weight during the treatment period. Data are presented as mean ± SEM. **(B)** Representative gross images of excised lesions from each experimental group collected at the study endpoint (day 10). Top labels indicate mouse number and side of plug (R:right, L:left), bottom numbers are the corresponding weight of the plug, in mg. Dashed boxes indicate the representative lesion with the median size for each group illustrated in Figure 5. Orange circles highlight the representative higher magnification lesions on the right. HUVEC TIE2-WT plugs are shown for comparison - they do not form highly vascularized lesions as mutant HUVEC TIE2-L914F. Scale bars, 1 cm. **(C)** Quantification of xenograft explant weight at day 10. All data points are presented in the violin plot. n=12 vehicle, 14 NSC treated, 6 TIE2-WT plugs in each group, from n=6,7,3 mice per group, respectively.

**Supplementary Table S1**: List of antibodies

| No. | Antibody / Reagent | Company | Catalog number | dilution |
|---|---|---|---|---|
| 1 | TIE2 (AB33) Mouse mAb | Cell Signaling | 4224S | 1:1000 |
| 2 | Phospho-TIE2 (Y992) | R&D Systems | AF2720 | 1:200 |
| 3 | RAC1 Antibody | Thermo Fisher | 1862341 | 1:1000 |
| 4 | GAPDH | Sigma/Millipore | MAB374 | 1:5000 |
| 5 | AKT | Cell Signaling | 2920S | 1:500 |
| 6 | Phospho-AKT (Ser473) | Cell Signaling | 4060T | 1:500 |
| 7 | ERK1/2 | Cell Signaling | 9101S | 1:500 |
| 8 | Phospho-ERK1/2 (Thr202/Tyr204) | Cell Signaling | 4696S | 1:500 |
| 9 | UEA-1 (Biotinylated) | Vector Laboratories | B-1065 | 1:200 |
| 10 | Rhodamine Phalloidin | Invitrogen/Fisher | R415 | 1X |

**Supplementary Figure S10**

### Full blots

### Figure 1C

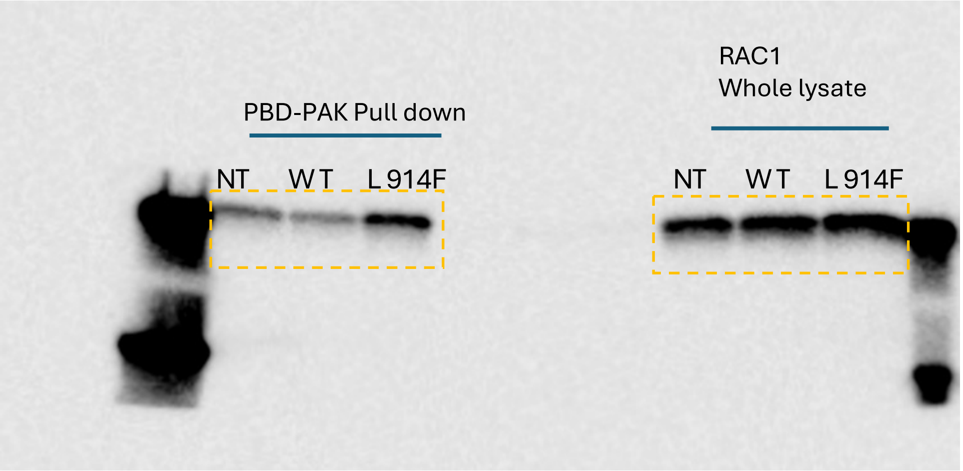

### Figure 4A

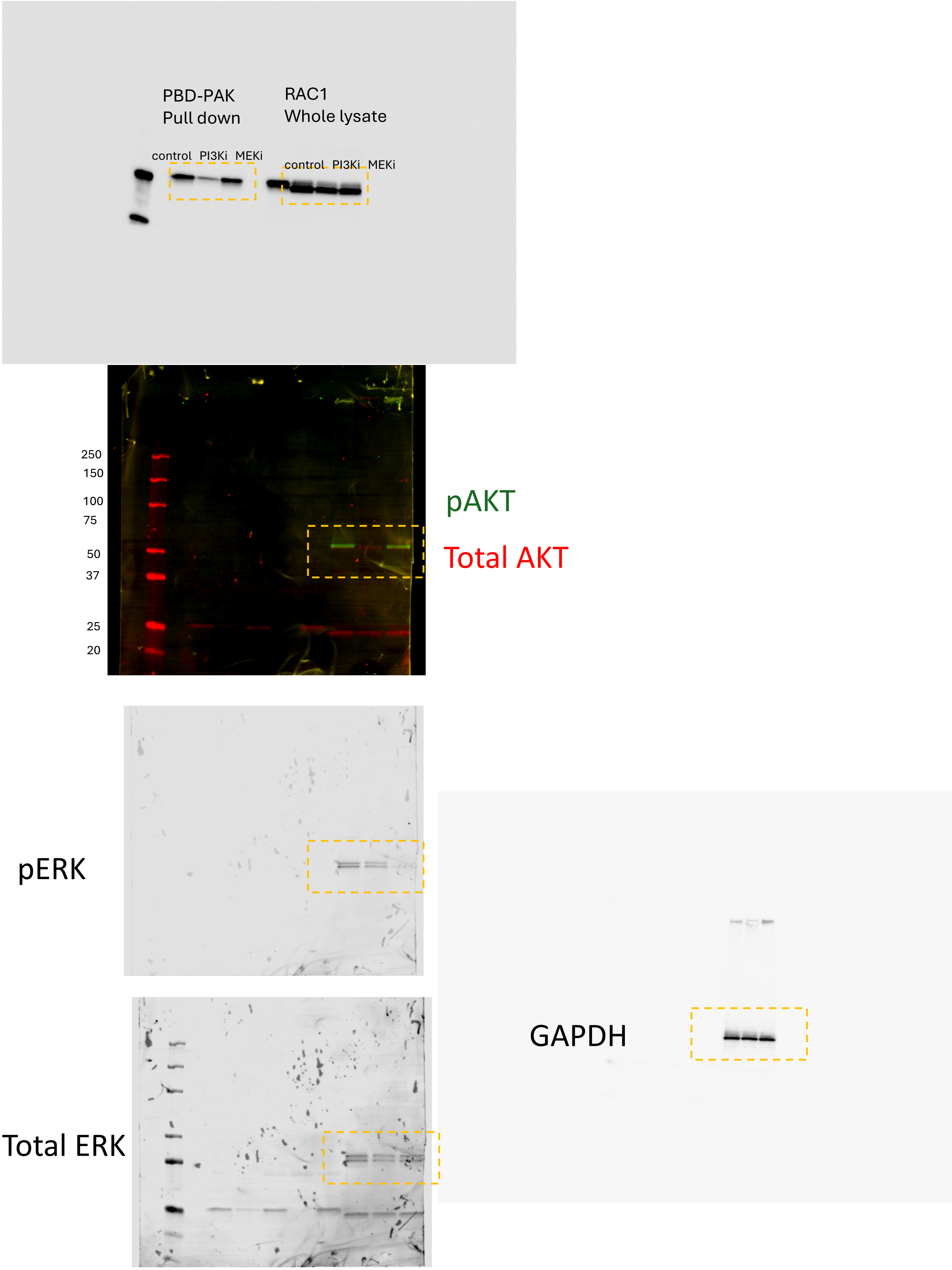
PBD-PAK
Pull down
RAC1
Whole lysate
control PI3Ki MEKi
control PI3Ki MEKi
250
150
100
75
50
37
25
20
pAKT
Total AKT
pERK
GAPDH
Total ERK

### Figure 4C

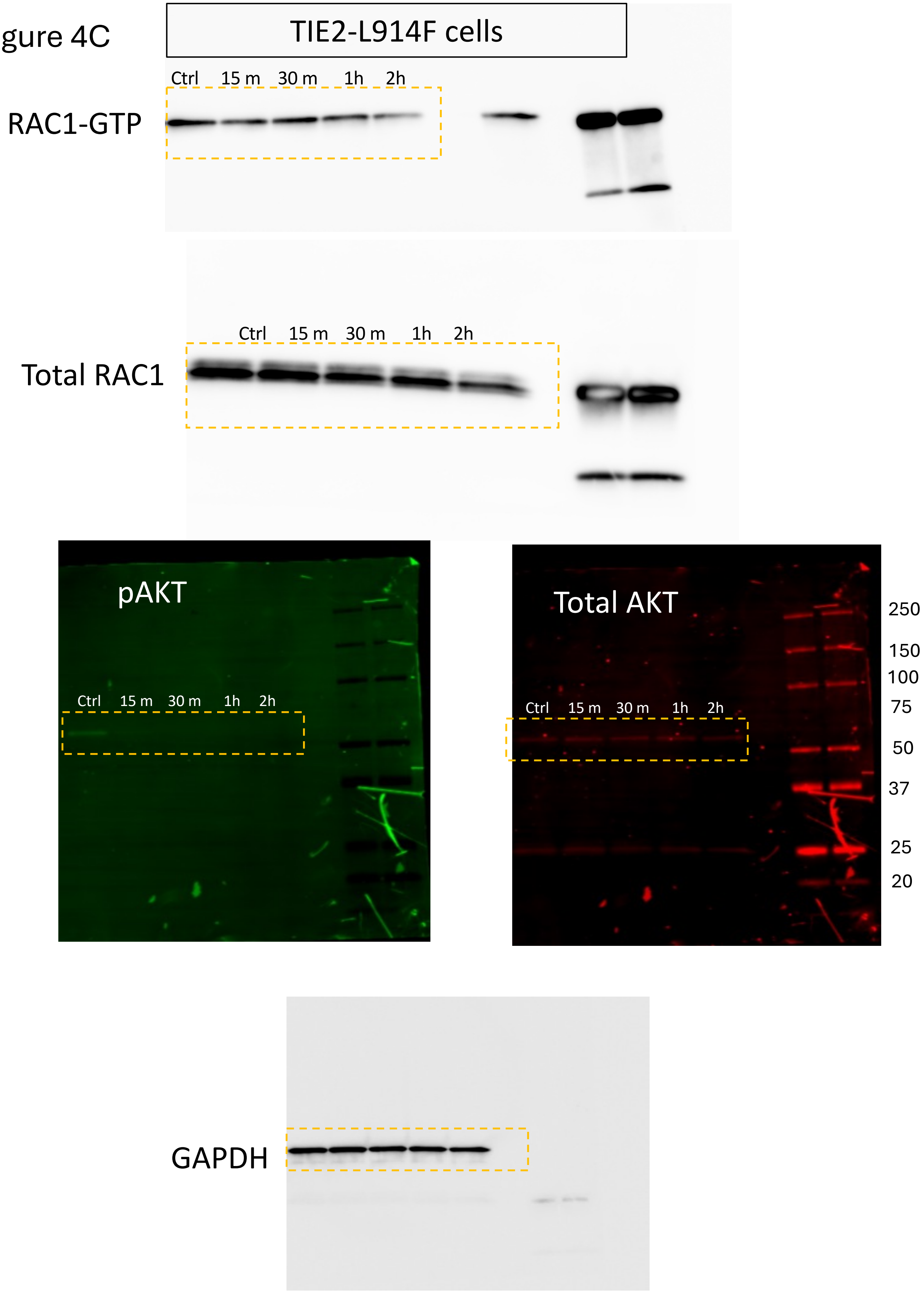
TIE2-L914F cells
Ctrl 15 m 30 m 1h 2h
RAC1-GTP
Ctrl 15 m 30 m 1h 2h
Total RAC1
pAKT
Ctrl 15 m 30 m 1h 2h
Total AKT
Ctrl 15 m 30 m 1h 2h
250
150
100
75
50
37
25
20
GAPDH

### Figure 4D, blots for quantification

TIE2-L914F cells

#### 1st experiment

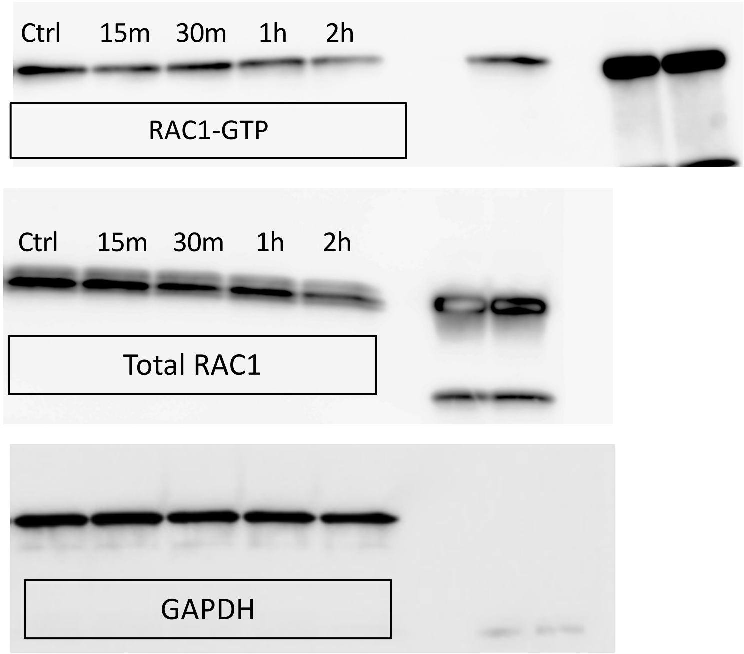


#### 2nd experiment

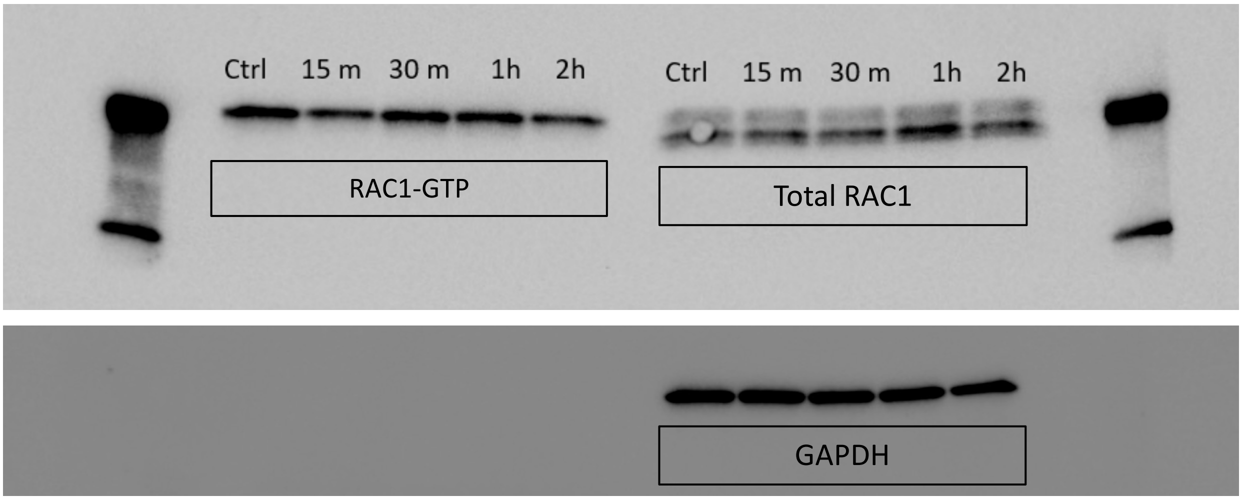


#### 3rd experiment

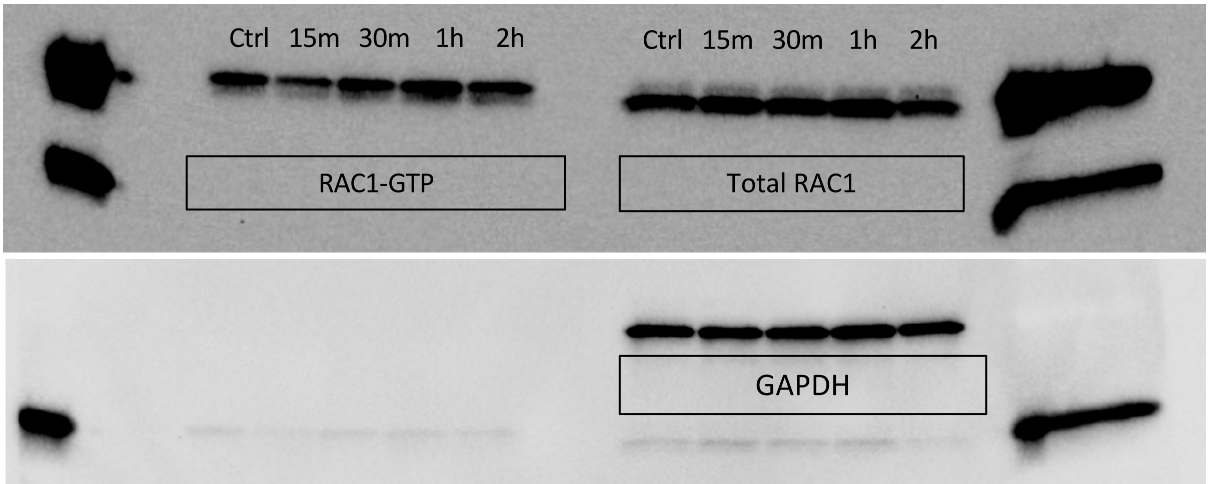

Supp Fig s1

pTIE2

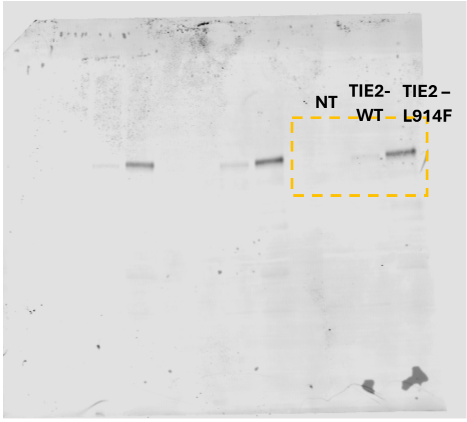


Total TIE2

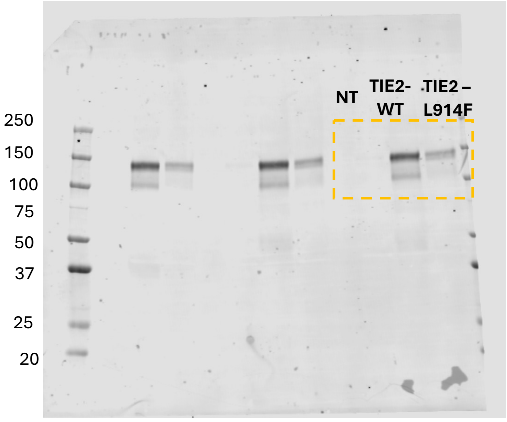


pAKT

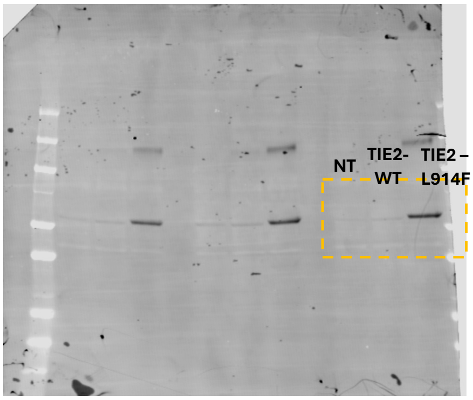


Total AKT

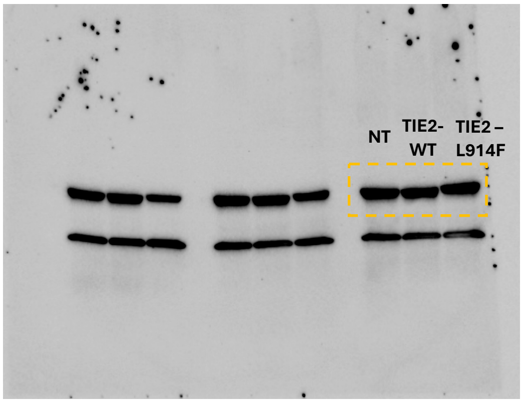

### Supp Fig s3

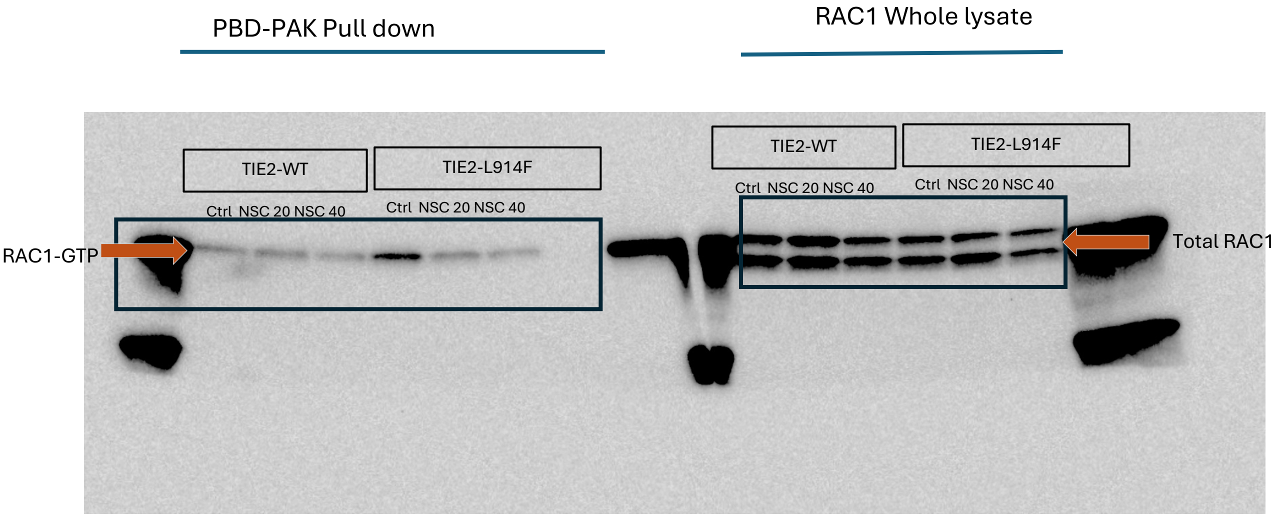
PBD-PAK Pull down
RAC1 Whole lysate
TIE2-WT
TIE2-L914F
TIE2-WT
TIE2-L914F
Ctrl NSC 20 NSC 40
Ctrl NSC 20 NSC 40
Ctrl NSC 20 NSC 40
Ctrl NSC 20 NSC 40
RAC1-GTP
Total RAC1

Supp Fig s6C

RAC1

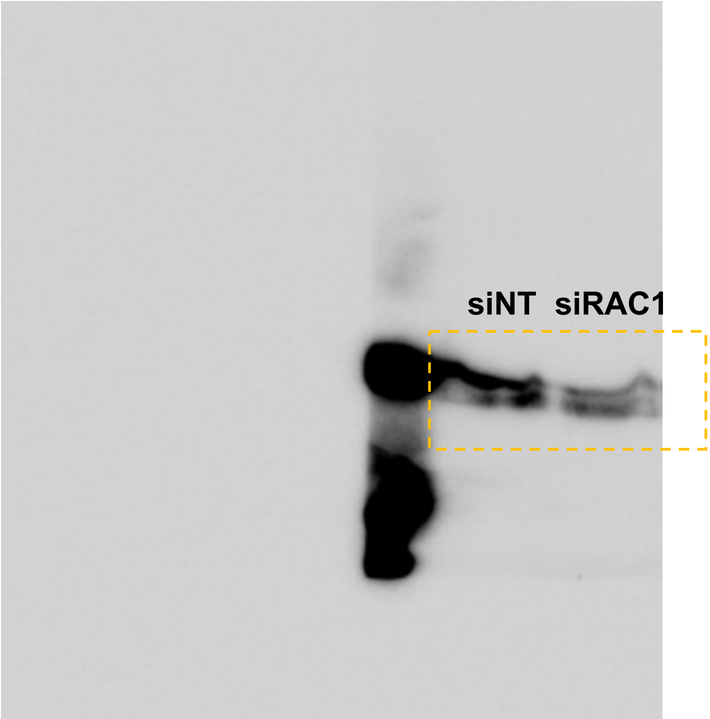


GAPDH

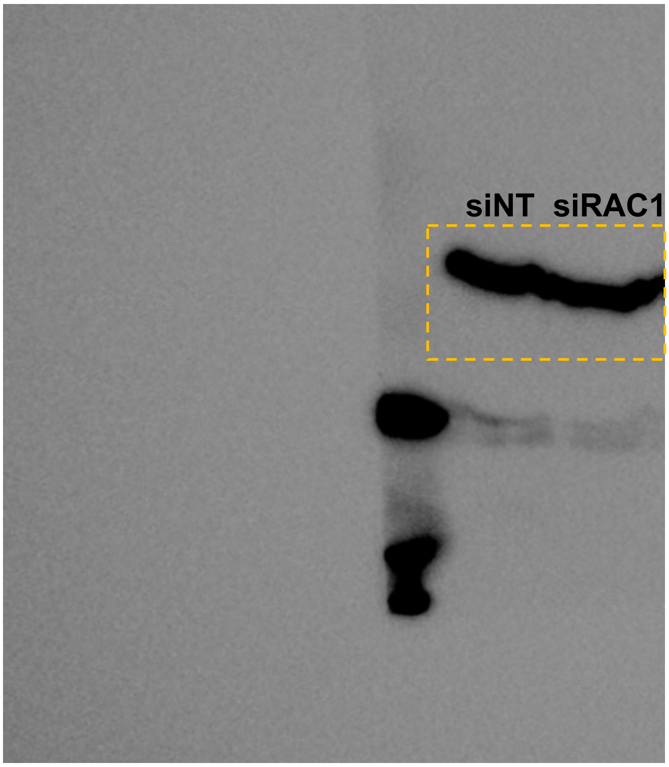

Supp Fig s8 A

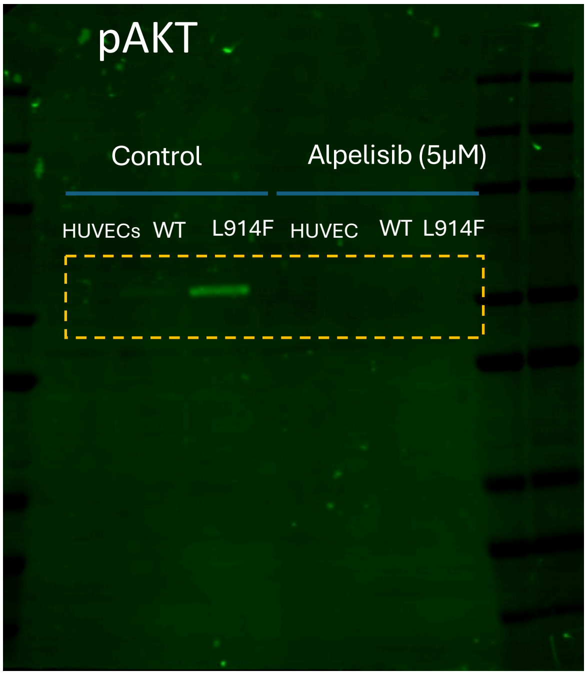


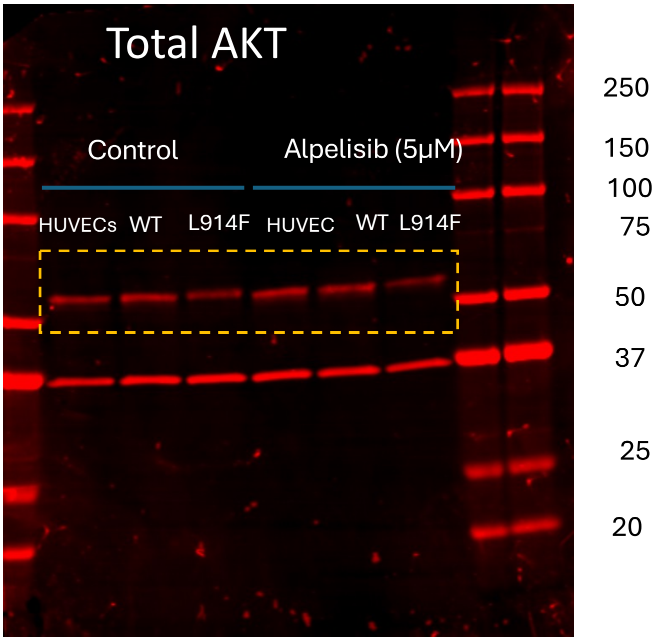


GAPDH

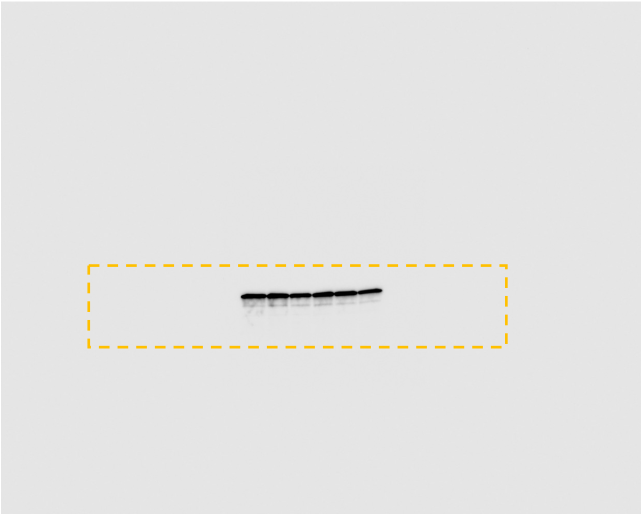

Supp Fig s8 B

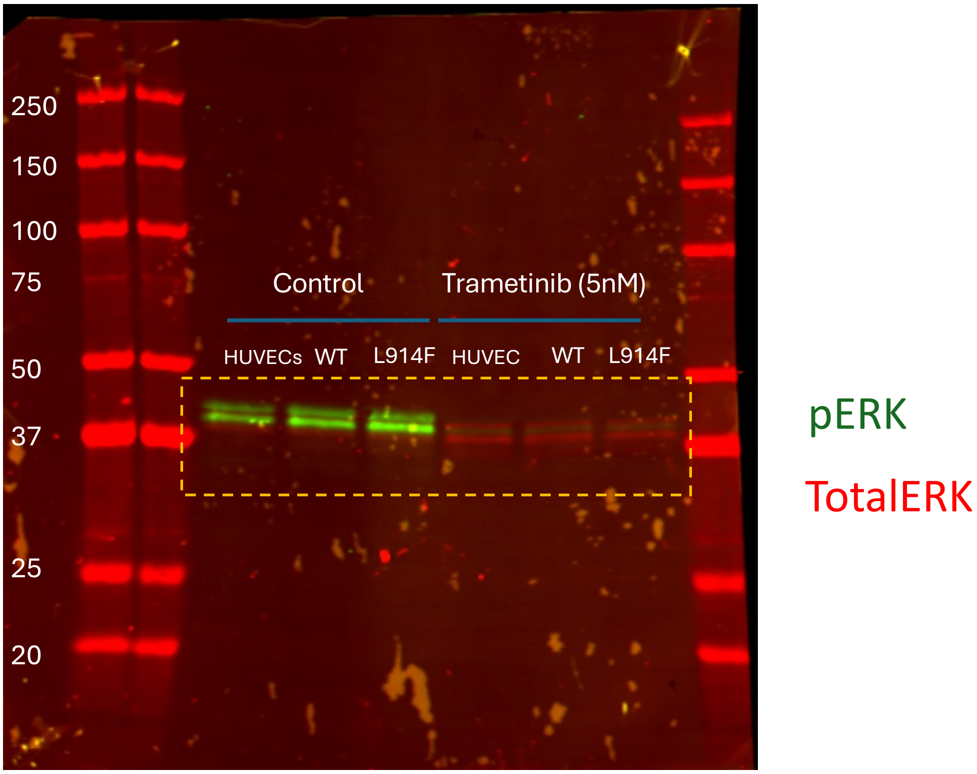
250
150
100
75
Control
Trametinib (5nM)
HUVECs WT L914F HUVEC WT L914F
50
37
25
20
pERK
TotalERK

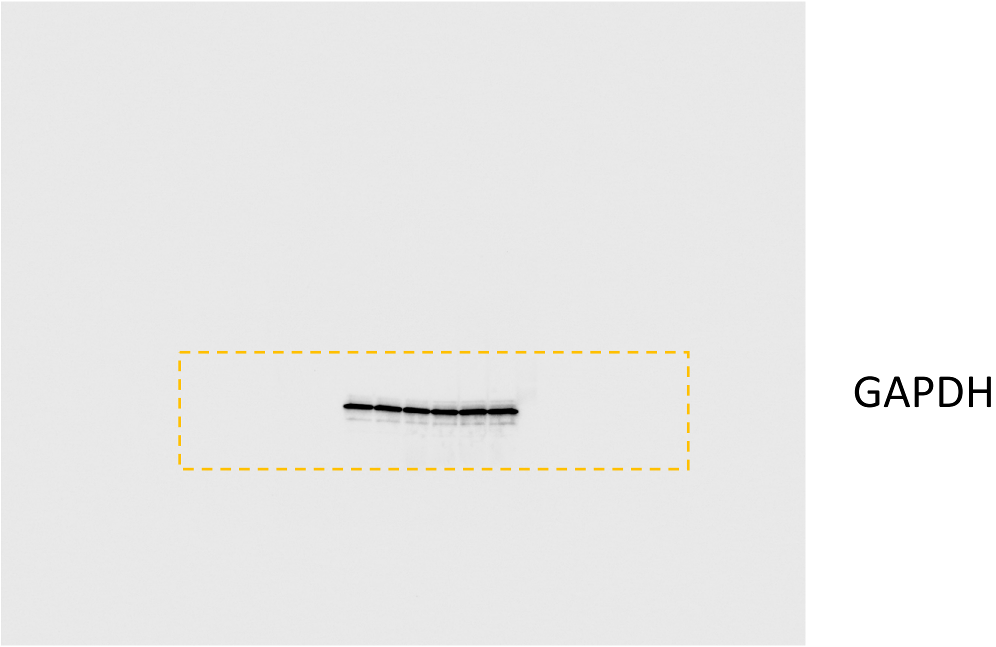
GAPDH
